# Niche differentiation is spatially and temporally regulated in the rhizosphere

**DOI:** 10.1101/611863

**Authors:** Erin E. Nuccio, Evan Starr, Ulas Karaoz, Eoin L. Brodie, Jizhong Zhou, Susannah Tringe, Rex R. Malmstrom, Tanja Woyke, Jillian F. Banfield, Mary K. Firestone, Jennifer Pett-Ridge

## Abstract

The rhizosphere is a hotspot for microbial C transformations, and the origin of root polysaccharides and polymeric carbohydrates that are important precursors to soil organic matter. However, the ecological mechanisms that underpin rhizosphere carbohydrate depolymerization are poorly understood. Using *Avena fatua*, a common annual grass, we analyzed time-resolved metatranscriptomes to compare microbial function in rhizosphere, detritusphere, and combined rhizosphere-detritusphere habitats. Population transcripts were binned with a unique reference database generated from soil isolate and single amplified genomes, metagenomes, and stable isotope probing metagenomes. While soil habitat significantly affected both community composition and overall gene expression, succession of microbial functions occurred at a faster time scale than compositional changes. Using hierarchical clustering of upregulated decomposition gene expression, we identified four distinct microbial guilds populated by taxa whose functional succession patterns suggest specialization for substrates provided by fresh growing roots, decaying root detritus, the combination of live and decaying root biomass, or aging root material. Carbohydrate depolymerization genes were consistently upregulated in the rhizosphere, and both taxonomic and functional diversity were high in the combined rhizosphere-detritusphere—suggesting coexistence of rhizosphere guilds is facilitated by niche differentiation. Metatranscriptome-defined guilds provide a framework to model rhizosphere succession and its consequences for soil carbon cycling.

## INTRODUCTION

The rhizosphere is a critical zone for C transformations in the terrestrial biosphere, since roots are the primary source of soil organic matter (1–3) and can significantly alter the rate of soil C turnover (4–6). Plants deposit a significant proportion of their photosynthates into soil as root biomass or exudates (7), and plant-derived polymeric carbohydrates such as cellulose and hemicellulose are the most abundant polysaccharides in soil (8, 9). These rhizodeposits create a high resource, high activity environment, and stimulate a bloom of microbial biomass (10) that undergoes ecological succession as roots grow and senesce (11, 12), selecting for organisms that benefit mineral nutrition (13) and overall plant health (14). Rhizodeposits also stimulate depolymerization by cellulases, chitinases, and proteases (15, 16) leading to higher rates of decomposition, and thus nutrient availability, in the region surrounding both living roots and decaying root detritus (17). However, the ecological controls of rhizosphere carbohydrate depolymerization are not well understood, which limits our ability to accurately model soil C dynamics (18) and plant-microbe interactions.

Previous studies suggest that rhizosphere community assembly is due to selective processes such as niche differentiation or habitat filtering (19–21), and metagenome sequences indicate the rhizosphere selects for microbial genomes with functional capacities that are distinct from bulk soil, where carbohydrate active enzyme (CAZy) genes are enriched in the rhizosphere (19, 21–23). However, it is unclear if this large genomic potential translates to high carbohydrate degradation activity in the environment. Genomic composition represents the full functional repertoire of a microorganism, the “fundamental metabolic niche” that constrains all the potential habitats it could hypothetically occupy (24, 25). But microbial communities contain functional redundancy that is not necessarily realized or expressed in the ecosystem (26). To understand “realized” metabolic niches within complex rapidly changing microbial communities (26), it is essential to consider expressed functional measurements— such as transcripts, proteins, or metabolites—that can reflect niche differentiation in real time (27, 28).

Measurements of expressed functions provide a useful way to study community assembly based on shared activities rather than shared phylogeny, and allow us to define microbial guilds—cohorts of organisms defined by similar function that is not dependent on phylogeny (29). In cases like the rhizosphere and detritusphere, where communities might logically be defined by functional traits rather than taxonomic relatedness, guilds defined by gene expression, rather than species, may be the most relevant parameter for understanding patterns of diversity (30), modeling community interactions (31), and identifying the gene transcripts that mediate root-accelerated decomposition. However, the ideal parameters for identifying or operationally defining guilds in microbial communities are unresolved. Microbial guilds have been identified previously by adding single substrates to soil and measuring subsequent increases in taxonomic relative abundance (32). We theorize that functional guilds can also be identified using population-resolved gene expression, where guild members turn on and off genes in a coherent spatial or temporal manner in response to the same habitat, resources, or environmental perturbations. Genome-centric analyses now allow us to track transcription in individual populations, which may be a more relevant approach than grouping transcripts across disparate classes or phyla (33).

Using comparative metatranscriptomics, we studied microbial degradation of macromolecular plant compounds, hypothesizing that gene expression would reflect distinct functional succession patterns in different soil habitats (rhizosphere and detritusphere), consistent with niche differentiation. The transcripts were extracted from soil near live and decaying roots in microcosms containing *Avena fatua*, a common annual grass, growing in its native soil. Population transcripts measured over the course of three weeks were binned using a genome-resolved reference database specific to our experimental soil. We found that carbohydrate depolymerization was executed by a series of microbial guilds, with distinct spatial and temporal response patterns in gene expression. We tested whether these guilds had differing life history traits based on their preferred substrate (rhizosphere or detritusphere), and assessed whether carbohydrate depolymerization expression was controlled by: a) increasing population size, b) upregulating transcription, or c) synergistically upregulating transcription in response to combined resources (i.e., combined rhizosphere-detritusphere). Our work provides a mechanistic framework for understanding the drivers of rhizosphere succession and identifies carbohydrate/lignolytic gene transcripts mediating root-accelerated decomposition.

## METHODS

### Experimental Design

The annual grass, wild oat (*Avena fatua*) was grown in two-chamber microcosms with a sidecar region designed to allow access to the rhizosphere (Fig. S1) (10, 16, 34). The outer wall of the sidecar was clear plastic, allowing us to monitor root growth and rhizosphere age. Microcosms were packed with soil (1.2 g/cm^3^) collected beneath a stand of *A. barbata* at the Hopland Research and Extension Center (Hopland, CA, USA). The soil is a Bearwallow-Hellman loam, pH 5.6 with 2% total C (35). For half the microcosms, 50 g soil was amended with 0.4 g dried *A. fatua* root detritus and spread on top of 100 g of sidecar soil; root detritus (also called ‘root litter’) had been grown in sand, triple washed, aged 1 year, and chopped to 1 mm. Each microcosm also contained a 1 μm mesh ‘bulk soil’ bag, designed to exclude roots but allow moisture equilibration; these contained 2 g soil, either amended with 0.016 g detritus (bulk + detritus) or unamended (bulk). Plants were grown in the main chamber for six weeks before starting the experiment. Six days prior, the divider separating the main chamber and sidecar was replaced with a slotted divider, and microcosms were tilted 40°, allowing roots to grow into the sidecar.

Root age was tracked in order to collect rhizosphere soil of defined age. New root growth was marked three days after the microcosms were tilted, and harvests took place 3, 6, 12 or 22 days later (Fig. S1). At each timepoint, we destructively sampled paired rhizosphere and bulk soil for two treatments (with and without detritus) with 3 biological replicates, collecting 48 total samples (24 rhizosphere, 24 bulk).

### Sample Collection

Rhizosphere soil <2 mm from the root was excised with a scalpel. Root sections and adhering soil were placed immediately in ice cold Lifeguard Soil Preservation Reagent (MoBio), vortexed for 2 min on medium speed, and pelleted according to the Lifeguard protocol. Roots were removed using flame-sterilized tweezers and supernatant removed. Pelleted soils were frozen on dry ice and stored at −80°C. Bulk soils were processed identically. Approximately 1 g of soil was collected per sample.

The remaining sidecar soil was collected for edaphic characterization. Soil pH was measured as per Fierer and Jackson (36) with a Corning 340 pH meter. Gravimetric moisture content was determined by measuring water loss from 10 g fresh soil after 2 days at 105°C. Total carbon (TC) was measured on a subset of the samples to calculate C addition due to the root material using an elemental analyzer IRMS (PDZ Europa, Limited, Crewe, UK).

### DNA/RNA Extraction

DNA and RNA were co-extracted from 0.5 g of frozen soil using a phenol-chloroform extraction protocol (37, 38). DNA and RNA were separated using the Qiagen AllPrep kit. RNA was treated with TURBO DNase (Thermo Fisher Scientific) following the manufacturer’s protocol and concentrated by ethanol precipitation. RNA was visualized an Experion Electrophoresis System (Bio-Rad), and quantified using the Qubit RNA BR Assay Kit (Thermo Fisher Scientific).

### Single Amplified Genomes

Rhizosphere and root endophyte microbial cells were sorted to create single amplified genomes (SAGs). Three grams of roots coated in rhizosphere soil were washed in 10ml of cell release buffer (0.5% Tween, 2.24 mM Na pyrophosphate in PBS), vortexed for 1 min at full speed on a horizontal shaker, and soft pelleted at 2KxG for 2 min (repeated 4 times). Cells from the supernatant were stained with SYBR, imaged using a Zeiss Axioimager M2 (CNR Biological Imaging Facility, UC Berkeley), and counted using ImageJ (39). Glycerol was added to the supernatant at a final concentration of 15% and frozen at −80° C prior to sorting. Roots were washed in PBS, shaken in 10% bleach (1 min), rinsed in water, dried by centrifugation, and frozen at −80° C prior to maceration using a sterile mortar and pestle prior to cell sorting. Thawed rhizosphere and root endophyte cells were isolated using fluorescence activated cell sorting, amplified by multiple displacement amplification, and screened using 16S rRNA sequencing (40).

### Sequencing Library Preparation

Metatranscriptomes, iTags (16S, ITS), and single amplified genomes (SAGs) were sequenced at the Joint Genome Institute (JGI); see Supplemental Methods for full details. Briefly, for metatranscriptomic libraries, ribosomal RNA was depleted using the Ribo-Zero rRNA Removal Kit (Epicentre) for Plants and Bacteria and reverse transcribed into cDNA. cDNA was sequenced (2×150bp) on an Illumina HiSeq2000 sequencer using a TruSeq SBS sequencing kit (v3). For iTag analysis, paired DNA and RNA were amplified from the same nucleic acid extract prepared for metatranscriptomics. iTag libraries targeted the bacterial 16S V4 region (primers 515F, 805R) (41, 42) and the fungal ITS2 region (primers ITS9, ITS4) (43, 44) using barcoded reverse primers (41). Amplicons were sequenced (2×300bp) on an Illumina MiSeq sequencing platform using a MiSeq Reagent Kit (v3 600 cycle). Selected SAGs that successfully amplified 16S rRNA were sequenced using the Illumina NextSeq platform (40).

### Sequence Processing

Metatranscriptomic raw reads were quality-trimmed (Q20) using fastqTrimmer, and artifacts were removed using DUK (45). Contaminating ribosomal RNA and transfer RNA were identified and removed with bowtie2 (46) by mapping reads against SILVA (47), Greengenes (48), IMG rRNA, GtRNAdb (49), and tRNADB-CE (50) databases. In total we sequenced 408 Gbp of RNA, and after *in silico* contaminant filtering, we obtained an average of 43 million paired-end metatranscriptomic reads per library (see Table S1 for repository IDs and sequencing statistics). We did not detect any bias towards rhizosphere or bulk soil in either sequencing library size (Figure S2) or gene diversity (Figure S3).

Amplicons were analyzed on JGI’s iTag analysis pipeline (iTagger) (41), which created OTUs at the 97% and 95% identity level for bacterial 16S and fungal ITS, respectively. Contaminants were removed using DUK, merged with FLASH (51), and dereplicated. Dereplicated sequences were sorted by decreasing abundance, clustered with USEARCH (52), and assigned taxonomy using the RDP classifier (53). SAG sequences were processed using BBTools (54). Sequences were filtered using BBDuk and mapped against masked contaminant references (human, cat, dog) using BBMap and BBMask. Reads with an average kmer depth <2 were removed. Normalization was performed with BBNorm and error correction with Tadpole (54). Sequences were assembled using SPAdes (version v3.7.1) (55); 200bp was trimmed from contig ends; contigs were discarded if length was < 2kbp or read coverage was < 2. Cross-contaminated contigs were identified and removed using CrossBlock (54). Automated SAG decontamination was performed with ProDeGe (version 2.3) (56) and assemblies were discarded if the total size was < 200 kbp. Details about the final draft assemblies are listed in Table S2.

### Soil-Specific Reference Database

Transcripts were mapped against a genome database specific to our Hopland CA experimental soil, composed of 96 metagenome-assembled genomes (MAGs) (NCBI PRJNA517182), 221 MAGs from a stable isotope probing (SIP) rhizosphere density gradient (57) (http://ggkbase.berkeley.edu/), 39 isolate genomes (58), and 29 single amplified genomes (SAGs) (this study; Table S1). Reference genomes were dereplicated using whole pairwise genome alignments at 98% nucleotide identity (59); we selected the highest quality representative based on completeness of single copy genes (60). To ensure we did not include multiple fragmented genomes from the same organism, genomes > 70% complete were clustered into groups that overlapped by at least 50%; genomes < 70% complete were clustered in a second round using a 30% overlap. The highest quality representative was selected for each cluster (score = # single copy genes – 2 * multiple single copy genes; the genome with the highest N50 was selected to break a tie). This resulted in 335 total genomes for our custom reference database (Table S3), composed of 214 rhizosphere SIP-MAGs (64%), 53 soil MAGs (16%), 39 isolate genomes (12%), and 29 SAGs (9%).

### Gene Annotation and Counts

Gene prediction was performed on all genome bins using Prodigal in metagenome mode (61). Protein sequences were annotated using dbCAN2 (62) (accessed April 2017), KEGG (63), and ggKbase (http://ggkbase.berkeley.edu/). Proteins with CAZyme functional domains were manually curated to generate a consensus annotation: CAZymes without KEGG or ggKbase annotations were ignored, and if the KEGG and ggKbase annotations disagreed, KEGG was selected. Genes containing signal peptide signatures for extracellular protein transport were annotated using SignalP 4.1 (64).

Trimmed and filtered reads were mapped against our soil-specific database using BBSplit (54). Reads that mapped ambiguously to multiple reference genomes were discarded to prevent double-counts. Transcripts were binned into population transcriptomes using a relaxed similarity cutoff (80% min identity) and should not be interpreted as genome transcriptomes. Gene counts were determined using featureCounts (R package: Rsubread).

### Data Analysis

Metatranscriptomic and amplicon sequencing data were normalized using DESeq2 to account for differences in sequencing effort (65), except for Shannon diversity analysis, where reads were rarified and diversity indices calculated using QIIME 1.9.1 (66). At each time point, significant differential expression relative to bulk soil was determined using DESeq2, which adjusts p values for multiple comparisons. Ordination and graph visualization were conducted in R (67). Data were ordinated using non-metric multidimensional scaling (R package: vegan), and significantly different clusters were determined using adonis (68). Correlations between environmental data and ordination data were tested using envfit (R package: vegan).

#### Carbohydrate Depolymerization CAZymes (d-CAZy)

We selected population transcriptomes with 4+ upregulated carbohydrate depolymerization genes for further analysis, which identified 26 of the 335 populations. Target substrates for depolymerization CAZymes (d-CAZy) were initially classified based on Berlemont and Martiny (69) and then refined into the following putative substrate categories using the consensus annotation described above: cellulose, xylan, xyloglucan, pectin, plant polysaccharides, microbial cell walls, starch and glycogen, xylose and cellulose oligosaccharides, oligosaccharides, mono- and disaccharides (see Table S7 for gene names and references). Area-proportional Venn diagrams were created to visualize the subset of the community that significantly upregulated d-CAZy transcripts relative to the total d-CAZy genomic potential (R package: venneuler).

#### Guild Assignment

We defined guilds based on d-CAZy expression over time and across treatments. Average d-CAZy differential gene expression (log2 fold change) relative to bulk soil was visualized using heatmaps (R package: pheatmap). One-dimensional hierarchical clustering was used to assign heatmap groups, which were classified as guilds based on resource preference and timing of peak gene expression. Maximum gene expression per enzyme per treatment was plotted by reference genome using ggplot2. Phylogeny is presented in accordance with current taxonomic nomenclature (70).

#### Decomposition Strategies

Decomposition strategies were assessed by evaluating gene expression levels relative to population abundance, and by comparing expression in the rhizosphere or detritusphere to the combined rhizosphere-detritusphere. Expression levels of the housekeeping genes Gyrase A and B were used as proxies for population abundance, as gyrases often have stable expression patterns over a variety of treatment conditions (71). Populations that increased in abundance, with significantly higher gyrase expression relative to bulk soil (by DESeq2), where increased d-CAZy expression is partially attributable to a larger population size were assigned to the ‘Grower’ strategy. Populations where d-CAZy expression was upregulated above population size (based on per capita gene expression), and d-CAZy fold-change was 3-fold higher than gyrase fold-change relative to bulk soil were assigned to the ‘Upregulator’ strategy. We use ‘Synergist’ to describe populations with 3-fold higher gene expression when combined resources were available (i.e., combined rhizosphere-detritusphere) compared to the rhizosphere or detritusphere alone.

## RESULTS

### Microcosm soil properties

Root detritus additions increased soil carbon from 2.0% ± 0.1 to 2.8% ± 0.1. Gravimetric soil moisture at the time of harvest averaged 0.34 ± 0.067 g water g^-1^ dry soil, with the exception of the final timepoint, when microcosms had an average of 0.11 ± 0.013 g water g^-1^ dry soil due to plant transpiration (Figure S4a). The addition of root detritus significantly increased bulk soil pH by 0.14 pH units at the first timepoint, and this differenced decreased over time (Figure S4b).

### Rapid community and functional assembly in the rhizosphere and detritusphere

Living roots and detritus rapidly altered bacterial community structure and functional assembly. All treatments diverged from bulk soil within 3 days, as seen by clear groupings in NMDS ordination space for both 16S cDNA and mRNA transcripts (Figure 1a and 1c, respectively). For 16S cDNA, the rhizosphere and detritusphere significantly shaped community composition (Figure 1) (see Table S4 for PERMANOVA F tables). Changes due to time explained 19% of the community variability, indicating that some taxonomic succession occurred within the treatments (Figure S5a). In contrast, fungal community composition, measured by ITS cDNA, was indistinguishable between rhizosphere and bulk soil (p > 0.1), and was instead significantly altered by both the detritus amendment and time (Figure 1b) (Table S4).

**Figure 1.**
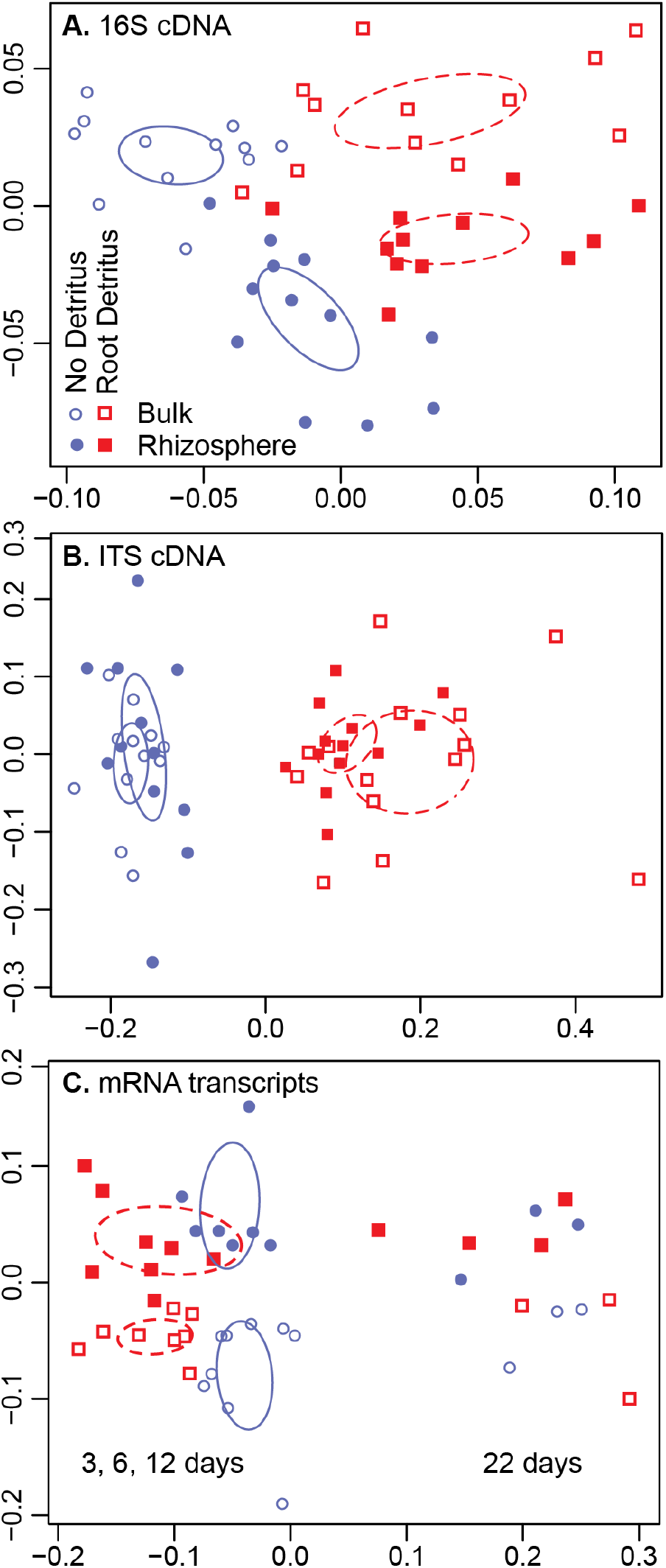
Influence of living roots and root litter on soil microbial communities and their gene expression during 3 weeks of *Avena fatua* root growth (independent harvests at 3, 6, 12, 22 days), as represented by NMDS ordination. Microbial community composition was measured by (A) bacterial 16S cDNA amplicons and (B) fungal ITS cDNA amplicons. Expressed functional composition was measured by (C) mRNA transcripts. Symbols represent four experimental habitats: rhizosphere (filled symbols), bulk soil (hollow symbols); each with added root detritus (red), or without added root detritus (blue). Ellipses represent the standard error of the weighted average of the centroid scores (calculated by ordiellipse). n=3 for each habitat and timepoint.

Time was the dominant factor structuring bacterial gene expression; transcripts from the final (22-day) timepoint clearly separate from earlier time points for all treatments (Figure 1c) (Table S4). This shift is correlated with both soil moisture (envfit: r^2^ = 0.87, p < 0.001) (Figure S5a) and time (envfit: r^2^ = 0.57, p < 0.001) (Figure S5b), likely because evapotransporation due to increased root biomass caused soil drying across all treatments at the final timepoint. When only the first three time points are considered (days 3, 6, 12), the rhizosphere and detritus treatments are the dominant factors structuring community gene expression (Table S4).

Early colonists of the rhizosphere rapidly increased in relative abundance within 3 days (16S cDNA, Table S5), and included Proteobacteria (Burkholderiaceae) and Verrucomicrobiota (Opitutaceae). Early colonists of the detritusphere included Fibrobacterota, Verrucomicrobiota (Chthoniobacteraceae, Opitutaceae), Armatimonadota, Bacteroidota, and Proteobacteria. In contrast, relatively few Actinobacteria and Acidobacteria significantly responded to either the rhizosphere or detritusphere on this timescale (3-22 days).

### Root detritus increased taxonomic and functional diversity

Root detritus amendment increased the taxonomic and functional Shannon diversity of both rhizosphere and bulk soil, and the combined rhizosphere-detritusphere had the highest overall taxonomic diversity by the final timepoint (Tukey HSD analysis, Figure 2a). Shannon taxonomic diversity was calculated based on 16S rRNA genes. Root detritus amendment of bulk soil significantly increased KEGG functional diversity, and appeared to have a similar effect on rhizosphere soil, although the trend was not significant at p < 0.05 (Tukey HSD analysis, Figure 2b). Of our four treatments, rhizosphere soils (with and without root detritus) had the highest expressed functional diversity after 22 days of root growth.

**Figure 2.**
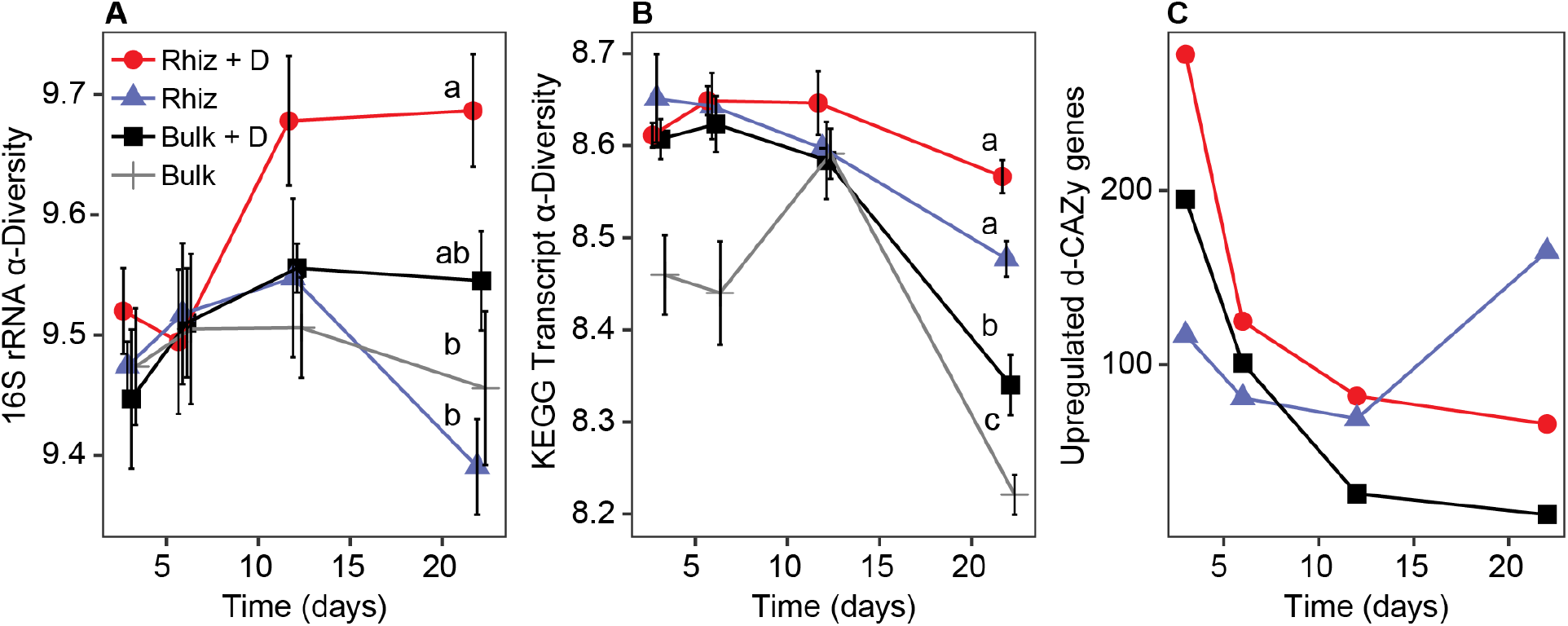
Taxonomic versus functional diversity in rhizosphere and bulk soils with and without detritus (+ D) harvested from *Avena fatua* microcosms over the course of 22 days. Average Shannon diversity for (A) 16S rRNA genes, and (B) KEGG functional genes derived from community mRNA transcripts. Error bars reflect one standard error. In order to make our results more comparable to a prior study of bacterial succession in the *Avena* rhizosphere, Shannon taxonomic diversity was calculated based on 16S rRNA genes (11). The different letters represent significant differences measured by Tukey HSD analysis at the final timepoint. (C) The cumulative number of significant differentially-upregulated decomposition CAZy (d-CAZy) genes relative to bulk soil, measured by DESeq. Treatments are: rhizosphere + detritus (red circle), rhizosphere (blue triangle), bulk soil + detritus (black square), and untreated bulk soil (grey cross).

### Roots stimulated expression of carbohydrate depolymerization transcripts

We curated a set of CAZyme genes relevant for plant and microbial carbohydrate depolymerization (d-CAZy, Table S7) and assessed expression of carbohydrate depolymerization transcripts relative to the bulk soil treatment. Overall, rhizosphere communities had the most significantly upregulated d-CAZy genes (Figure 2c). The combined rhizosphere-detritusphere had the largest number of significantly upregulated d-CAZy genes, with the exception of the final time point, when unamended rhizosphere soil had the largest number of upregulated genes. This result was generally consistent across the four major CAZyme classes (auxiliary activity, carbohydrate esterases, glycoside hydrolases, polysaccharide lyases) (Figure S6). In the bulk soil, root detritus additions initially stimulated a large pulse of d-CAZy activity but this dropped dramatically over time; by the final timepoint, only 10-20% of the genes were distinguishable from bulk soil expression levels.

### Realized niches in rhizosphere and detritusphere

The fundamental metabolic niche describes the full metabolic repertoire of a microorganism, and is represented by its total genomic content. For the individual populations in our reference database, we identified populations with statistically significant gene expression and compared it to genomic content in an effort to identify the ‘realized’ metabolic niches within our bacterial community (Figure 3). While many populations had the genomic capacity for carbohydrate depolymerization, only a small fraction significantly upregulated these genes relative to bulk soil by DESeq2. Populations upregulating cellulases and xylanases relative to bulk soil across the three treatments were 11-15% and 10-19% of the total genomic potential, respectively (Figure 3). The relative expression patterns for gyrase A and B housekeeping genes indicate that the general population dynamics followed similar patterns by treatment as observed for functional genes.

**Figure 3.**
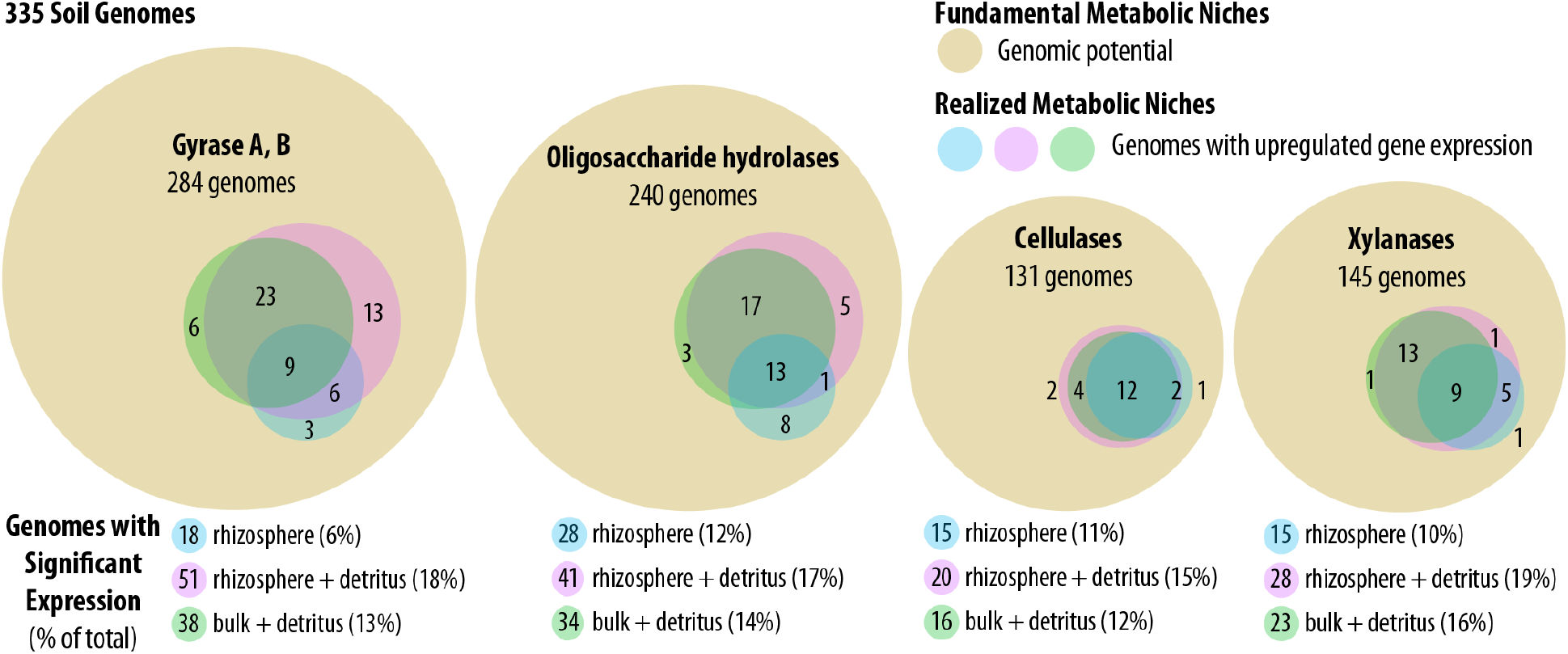
Fundamental (genomic potential) versus realized metabolic niches (upregulated gene expression) for key carbohydrate degradation gene classes. Area-proportional Venn diagrams indicate the number of functionally active taxa by soil habitat relative to the total metagenomic capacity for 335 assembled soil genomes. The outer circle (brown) indicates the number of unique genomes in the reference database with the genomic potential for the specified class of genes; inner circles reflect the number of taxa that differentially upregulated each class of genes relative to bulk soil for each treatment: rhizosphere (blue), rhizosphere + detritus (pink), and bulk soil + detritus (green). Overlapping regions represent shared niche space, with the number of genomes shared between different treatments. Genome classes analyzed include: gyrase A, B (housekeeping gene), oligosaccharide hydrolases (e.g., glycosidases, xylanases), cellulases, and xylanases (see Table S7 for full gene list). The bottom panel lists the number of active genomes by treatment, and the percentage of active genomes relative to total genomic potential is denoted in parentheses.

### Guilds defined by temporal and habitat gene expression

We assessed population gene transcription patterns over time and across habitats to define ecological guilds. The majority of differential gene expression came from 26 of our 335 reference genomes; these had at least four d-CAZy genes significantly upregulated relative to bulk soil. We note that 20 of them were derived from our rhizosphere SIP-metagenome database (Figure 4). These bacterial populations had distinct rhizosphere versus detritusphere transcriptional preferences (one dimensional hierarchical clustering, Figure 4). We averaged d-CAZy differential expression per population to show broad differential expression patterns (Figure S7). Using statistical upregulation of depolymerization genes as a proxy for resource preference and guild membership, we defined “Rhizosphere,” “Detritusphere,” and “Aging Root” guilds, and a “Low Response” group where there was no discernable habitat preference.

**Figure 4.**
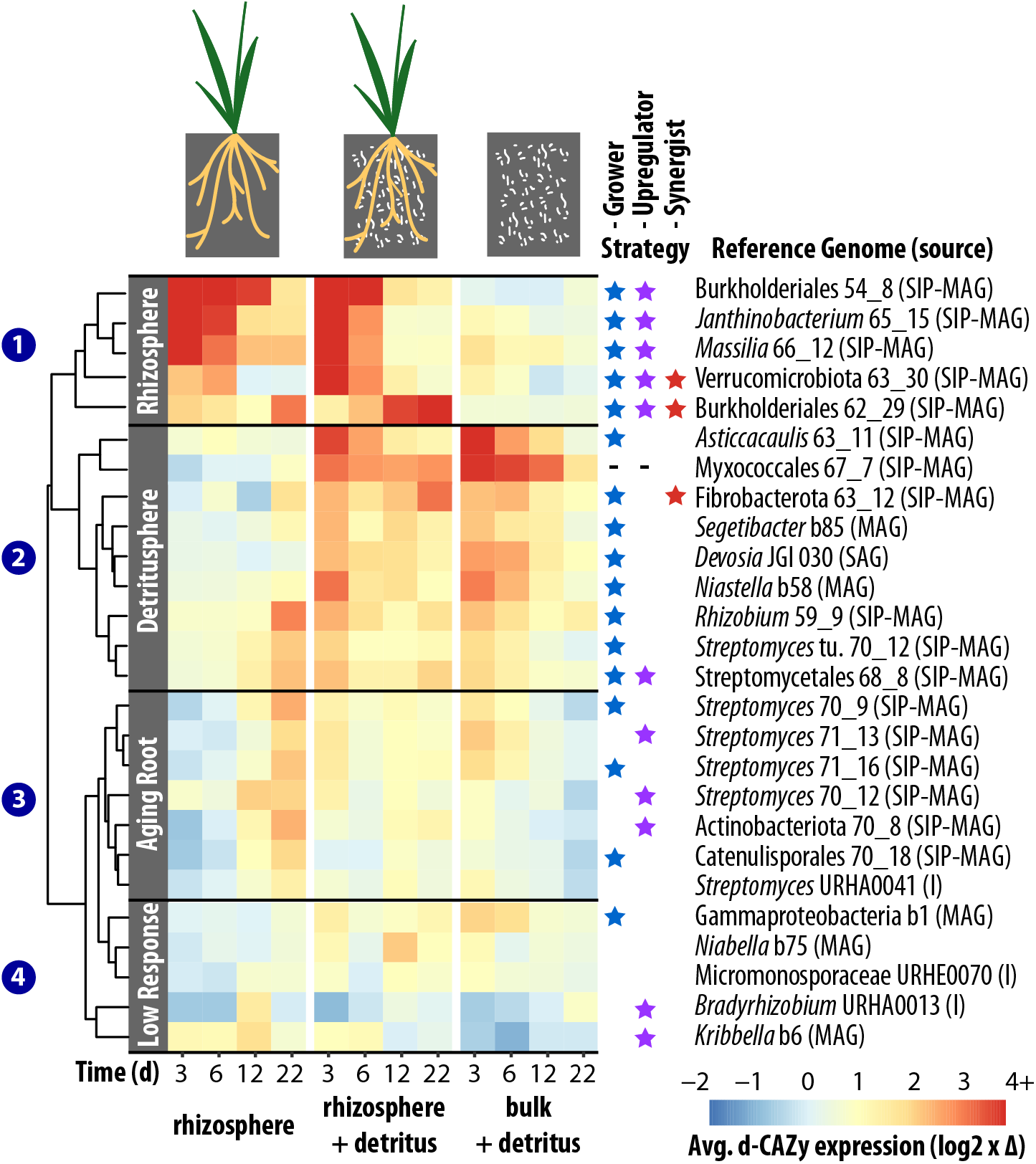
Time-series heatmap representing average decomposition CAZy (d-CAZy) gene expression per genome for 26 d-CAZy-responsive taxa during a 22-day *Avena fatua* microcosm experiment; responsive taxa significantly upregulate 4+ d-CAZy genes relative to bulk soil. Red indicates log2-fold gene upregulation in the treatment, blue indicates gene upregulation in bulk soil. Reference genome taxonomy is listed for the population transcriptomes (rows), as is the source of the genome: rhizosphere SIP-metagenome (SIP-MG), soil metagenome (MG), cultured isolate genome (I), single amplified genome (SAG). Time (days) is indicated by the columns. Metatranscriptomic guild assignment was accomplished through one-dimensional hierarchical clustering and is denoted by the left gray bars and numbers; high d-CAZy gene expression when living roots were present were assigned to the ‘Rhizosphere’ guild; high d-CAZy expression when added detritus was present formed the ‘Detritusphere’ guild; high d-CAZy expression when living roots were present, but where expression peaked at the last timepoint, formed the ‘Aging Root’ guild. Stars indicate decomposition strategy: blue stars indicate populations that significantly increased abundance (‘Growers,’ Fig. S8)); purple stars indicate populations where per capita gene expression was 3x > abundance (‘Upregulators,’ Fig. S9)); red stars indicate ‘synergist’ populations, where gene expression in combined rhizosphere-detritusphere was 3x > than rhizosphere or detritusphere alone (‘Synergist,’ Fig. S10)). Hyphens indicate no gyrase data was available for the calculation.

### Carbohydrate depolymerization guilds undergo functional succession

Transcriptionally-defined guilds captured a functional succession in carbohydrate depolymerization, for both polysaccharides and also oligosaccharide breakdown products. The Rhizosphere and Detritusphere guilds had high d-CAZy expression within the first 6 days, then between 12-22 days an additional Aging Root guild emerged (Figure 4).

The Rhizosphere guild contained Proteobacteria order Burkholderiales and a Verrucomicrobiota population from the Opitutaceae (Figure 4, Group 1). Cellulases (endoglucanases), xylanases, and xyloglucanases were most highly expressed at 3 days, as were enzymes for potential breakdown products like cellulose- and xylan-oligosaccharide hydrolases (beta-glucosidases and beta-xylosidases, respectively) (Figure S7a-d). One Burkholderiaceae population did not follow this pattern, and instead had high d-CAZy expression at the final timepoint. Overall, xyloglucan hydrolases were characteristic of rhizosphere populations, and observed only once in the detritusphere (bulk + detritus) (Figure 5).

**Figure 5.**
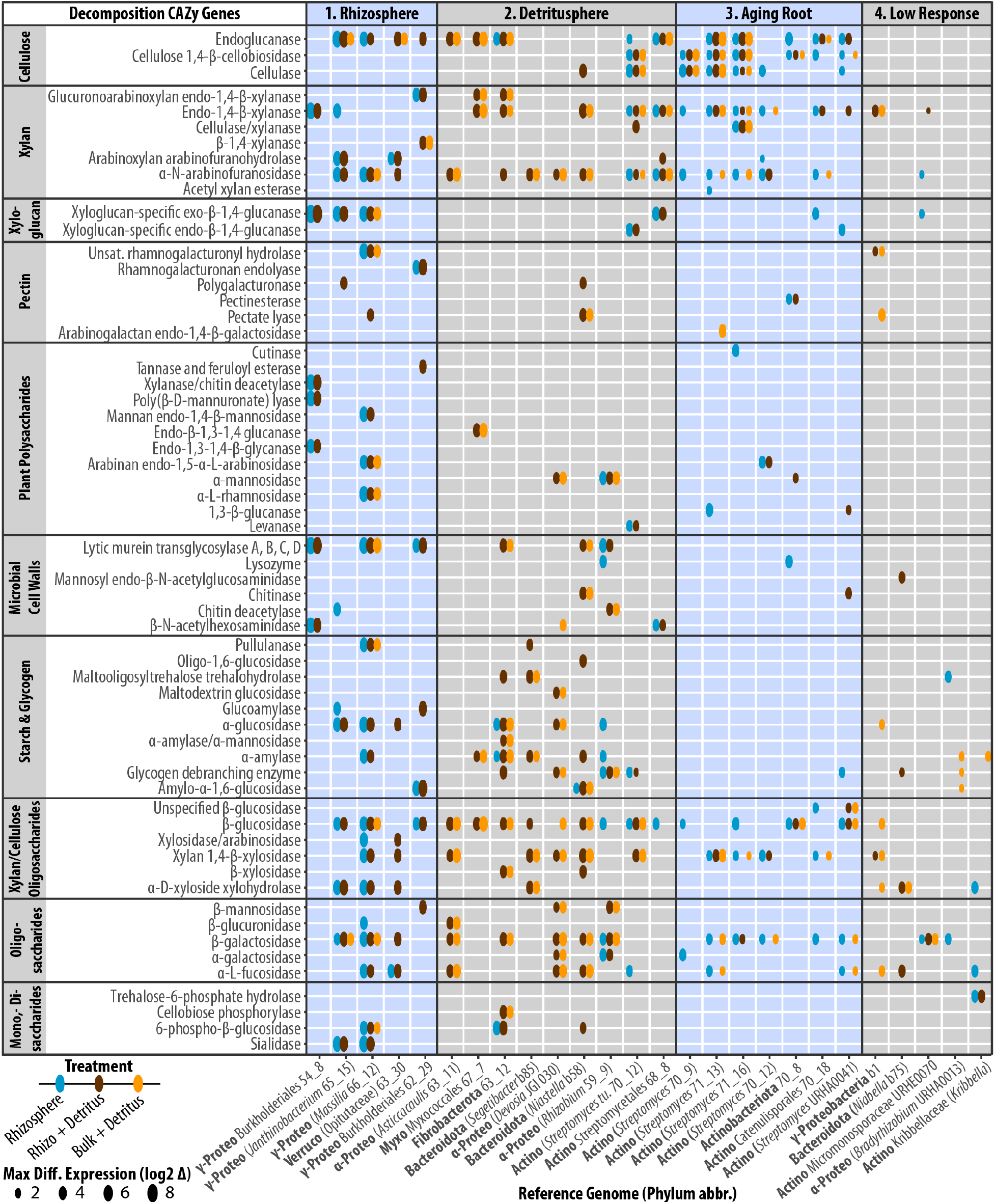
Upregulated decomposition CAZy genes for 26 bacteria classified into decomposition guilds defined in this study (Rhizosphere, Detritusphere, Aging Root, Low Response; see Fig. 4). Ovals and their size indicate maximum differential expression relative to bulk soil (log2-fold change) over the time course for the treatments: rhizosphere (blue ovals), rhizosphere + detritus (brown ovals), and bulk + detritus (yellow ovals). Genes are grouped by the enzyme’s putative target substrate: plant polysaccharides (cellulose, xylan, xyloglucan, pectin, other plant polysaccharides), microbial cell walls, starch and glycogen, xylan- and/or cellulose-oligosaccharides, other oligosaccharides, and mono- and disaccharides. Phylum abbreviations: *γ*-Proteobacteria (*γ*-Proteo), Verrucomicrobiota (Verruco), α-Proteobacteria (γ-Proteo), Myxococcota (Myxo), Actinobacteriota (Actino).

The Detritusphere guild was phylogenetically diverse, including members from Proteobacteria, Myxococcota, Fibrobacterota, Bacteroidota, and Actinobacteroita phyla (Figure 4, Group 2). With the exception of the Rhizobiaceae population, members of the Detritusphere guild typically upregulated cellulases and xylanase (or both) soon after detritus was added (3, 6 days), and cellulose- or xylan-oligosaccharide hydrolases for potential breakdown products (Figure 5).

In the Aging Root guild, Actinobacteriota populations from the *Streptomycetaceae* and Catenulisporales had high d-CAZy gene expression at the final timepoint (22 days) in the rhizosphere, and early gene expression in the detritus-amended treatments (Figure 4, Group 3). The Aging Root guild had almost no upregulated genes for starch, glycogen, cell wall, and disaccharide decomposition (Figure 5).

### Guild-based assessment of decomposition strategies

We used metatranscriptomic expression patterns to determine the prevalence of three decomposition strategies within the microbial guilds (Figure 4): a.) increased population abundance, b.) upregulated gene expression (above per capita abundance), or c.) synergistic gene upregulation when combined resources were available (i.e., combined rhizosphere-detritusphere). We interpreted significant gyrase upregulation relative to bulk soil as a population increase (DESeq2, Figure S8).

Decomposition strategies varied by guild membership, and were not mutually exclusive. All members of the Rhizosphere and Detritusphere guilds were “Growers” and increased in abundance (Figures 4 and S8). Rhizosphere guild populations were also “Upregulators” (3-fold higher gene expression per capita) (Figures 4 and S9). In contrast, enzyme expression in the Detritusphere guild tracked population levels; only one population was an “Upregulator”. Over half of the Aging Root guild did not change in abundance relative to bulk soil. This guild, composed entirely of Actinobacteriota and two *Steptomyces* populations, had “Upregulators” with populations sizes statistically indistinguishable from bulk soil.

Three “Synergist” populations upregulated gene expression in response to combined resources, with 3-fold higher gene expression in the combined rhizosphere-detritusphere compared to either habitat alone (Figures 4 and S10). These include Verrucomicrobiota (Opitutaceae) and Burkholderiales populations within the Rhizosphere guild, and a Fibrobacterota population from the Detritusphere guild. The 30S ribosomal protein S3 (RP-S3) from the Opitutaceae reference genome was 93% similar to the *Opitutus terrae* RP-S3 (by blastx (72)), an obligate anaerobe isolated from rice paddy soil. The Burkholderiales and Fibrobacterota reference genomes were most closely related to uncultivated MAGs (RP-S3 85% similar to Rhizobacter MAG and 73% similar to Fibrobacterota MAG, respectively). Both the Optitutaceae and Fibrobacterota reference genomes contained CBB3 cytochrome oxidases (putatively, a microaerophilic version of cytochrome oxidase); these genes were actively expressed but not upregulated relative to bulk soil. At the early timepoints, the Opitutaceae population upregulated enzymes for xylan degradation (arabinoxylan arabinofuranosidase) and xylan breakdown products (xylan 1,4-beta xylosidase, alpha-D-xyloside xylohydrolase) (Figures 5 and S7d). The Fibrobacterota population synergistically upregulated endoglucanases, endo-1,4-beta-xylanases, and enzymes targeting their potential breakdown products (beta-xylosidase, cellobiose phosphorylase) (Figure S7h). At later timepoints, the Burkholderiales upregulated putative lignocellulosic enzymes such as endoglucanase, tannase and feruloyl esterase, and rhamnogalacturonan lyase (Figure S7e).

## DISCUSSION

### Rapid community and functional assembly in the nascent rhizosphere and detritusphere

The soil microbial community surrounding roots undergoes a compositional succession corresponding to the phenological stages of plant growth (11, 12). However little is known about microbial gene expression during rhizosphere succession, and the temporal relationship between functional succession versus community changes. We used genome-centric, comparative metatranscriptomics to identify taxa mediating root-enhanced decomposition using carbohydrate gene transcripts. Since many soil taxa are non-cultivable by conventional methods, this approach offers insights into the physiologies of populations only known by sequencing (33). Our results, some of the first using a genome-centric metatranscriptome approach in soil, illustrate that different microbial populations have specialized functions and life strategies based on spatiotemporal differences in root habitats.

We found community and functional assembly proceeded at different rates—while taxonomic composition underwent minor successional changes over three weeks, expressed functional composition distinctly shifted between 12-22 days. mRNA has a short half-life relative to DNA, and is a sensitive indicator about ongoing ecological processes and near-real-time conditions experienced by cells (73). Our previous work indicates rhizosphere community composition continues to shift from three weeks until senescence (11), but the faster changes we observe for transcript inventories suggest microbes experience changes in rhizodeposits, environmental conditions (e.g. moisture, pH, O2), or other signals on the scale of days (77). The relative speed of functional shifts suggests that expressed functional succession occurs at a faster time scale than compositional changes, and possibly dictates the rhizosphere microbial community succession that occurs over longer time scales. This illustrates a benefit of using time-resolved metatranscriptomics to assess activity of specific microbial populations and the processes that lead to community assembly, since organisms transcriptionally respond to stimuli on a shorter time scale than evinced by replication.

### SIP-metagenomes produced the most useful genomes for soil metatranscriptomics

The proportion of population-specific transcripts mapping to our reference genomes illustrates the comparative benefits of the four sequence products in our custom database: single amplified genomes, isolate genomes, deeply sequenced bulk soil metagenomes, and rhizosphere stable isotope probing (SIP) metagenomes. Genomes derived from rhizosphere SIP-metagenomes proved to be the most relevant for transcript mapping, and the source of most of the populations with 4+ upregulated carbohydrate depolymerization genes. In a previous study (57), these populations also showed high ^13^C-incorporation by rhizosphere ^13^C-SIP (unpublished data), where the plants were continuously-labeled with ^13^CO2 for 6 weeks. Our results indicate that SIP-metagenome datasets may be a highly fruitful genomic resource for environmental metatranscriptomics and other omics analyses.

### Metatranscriptomic guilds provide a framework to understand rhizosphere succession

By assigning expressed carbohydrate depolymerization genes to individual population genomes derived from our custom genome database, we stepped beyond gene-centric studies that have shown rhizosphere gene expression with plant development (12) or environmental changes (74) and identified specific carbohydrate depolymerization guilds based on shared spatiotemporal gene expression.

We used these guilds to evaluate decomposition strategies that underpin altered carbohydrate degradation rates commonly found surrounding roots (4). In macroecology, the guild concept is a common way to group populations as functional ecological units, based on their resource utilization traits or life history strategies (25, 29). In environmental microbiology, while next-generation sequencing allows us to group microbial communities taxonomically (75), taxonomy and function may not correspond (26, 30, 76). We identified four guilds based on spatiotemporal CAZy expression patterns (Rhizosphere, Detritusphere, Aging Root and Low Response). This ecological categorization framework may be particularly useful for phylogenetically ubiquitous microbial functions—for example, soil organic matter decomposition (32, 76–78), stress, or nitrogen mineralization—where guilds are based on shared life history traits rather than phylogeny.

Within each guild, many populations engaged multiple catabolic pathways for carbohydrate degradation, including potential degradation by-products such as cellulose- and xylose-oligosaccharides. For example, populations from the Rhizosphere and Detritusphere guilds not only expressed enzymes for cellulose and xylan degradation, but also their breakdown products. Recent work suggests that facilitative processes such as cross-feeding in large networks can act to stabilize coexisting competitors for resources (79–81). Genome-resolved metagenomic analyses indicate the importance of metabolic byproduct handoff in linking together interacting members of microbial communities (80, 81). The breadth of carbohydrate degradation pathways that Rhizosphere and Detritusphere guilds engage in may be a potential explanation for the stable, positive and repeatable interaction networks in the rhizosphere that we observed in a previous study (82). We hypothesize that complex cross-feedings networks promote coexistence within highly interconnected rhizosphere communities (79).

### Niche differentiation promotes coexistence of rhizosphere and detritusphere guilds

By combining taxonomy and function, genome-resolved metatranscriptomics, we examined both the ‘potential’ and ‘realized’ metabolic niches (26, 33) of bacteria in our experiment. The niche differentiation concept asserts that organisms coexist by subdividing available resources, such as food or space (25, 32). As the number of niches increases in a system, so should the number of coexisting species (83). During root colonization in the combined rhizosphere-detritusphere, coexisting guilds began to develop, demonstrating that carbohydrate depolymerization preferences that were evident when root habitats were presented in isolation (rhizosphere or detritusphere), could coexist when combined. When detritus was added to the rhizosphere, the most populations predominantly demonstrated spatial and temporal coexistence rather than synergistic consumption of resources. These results are reflected by the higher functional alpha diversity in the combined rhizosphere-detritusphere; we saw approximately additive increases in functional diversity when a new resource (root detritus) was added to the system. Our work suggests that spatial and temporal niche differentiation promotes microbial coexistence in the rhizosphere and detritusphere.

### Guild-based assessment of carbohydrate degradation strategies

We further assessed if our guilds had differing d-CAZy transcriptional strategies and evaluated if our increases in gene expression (a) tracked increases in population size (‘Growers’), (b) were upregulated per capita (‘Upregulators’), or (c) synergistically upregulated when both root exudates and detritus were available (‘Synergists’). These strategies were not mutually exclusive, and their prevalence varied according to guild membership. All organisms in the Rhizosphere guild were both ‘Growers’ and ‘Upregulators,’ while the Detritusphere guild were primarily ‘Growers.’ Multiple studies have shown that the input of organic-C substrates can increase or decrease the rates of C degradation of surrounding soil organic matter, which is a phenomenon known as priming (4, 5, 84). Due to the large number of significantly upregulated decomposition transcripts in the rhizosphere, both with and without detritus amendments, this system has a high potential for increased rates of decomposition in the rhizosphere, as was previously observed in this plant-soil system (6). This is consistent with the expectations for positive rhizosphere priming, where fresh organic matter provided by the rhizosphere stimulates the production of enzymes that can degrade soil organic matter (17, 85, 86).

Based on the rhizosphere priming hypothesis, we would expect to observe “Synergists” in the combined rhizosphere-detritusphere. Strikingly, we only observed 3 “Synergist” populations, and two of these were putative microaerophiles. This suggests that these populations may also be partitioning their niches based on changes in the edaphic environment, such as oxygen or pH, rather than simply by consuming combined resources alone. The Verrucomicrobiota MAG is distantly related to Opitutaceae isolates derived from oxygen-limited rice patties and insect guts (87–90). Fibrobacteres include cellulose degrading bacteria found in mammal rumens (91), termite guts (92), anaerobic cellulose reactors (93), and rice paddy soil (94). Both MAGs contain cytochrome oxidases with high oxygen affinity (CBB3), which is associated with organisms living in microaerophilic environments (95). High amounts of heterotrophic respiration can create microaerophilic zones in otherwise aerobic environments, such as the rhizosphere (96, 97). The combined oxygen demand from both the rhizosphere and detritusphere may have been sufficiently high to create microaerophilic niche for root detritus decomposition, thus providing a possible mechanism for the observed synergistic response. Both of these populations are rhizosphere inhabitants found in our other studies (11, 20) suggesting that this synergistic decomposition in the combined rhizosphere-detritusphere may be functionally significant in semiarid grasslands.

During the functional succession of guilds, one guild emerged during the latter half of the experiment as the rhizosphere aged. Interestingly, more than half of the Aging Root guild had population sizes that were indistinguishable from bulk soil based on gyrase housekeeping gene expression, but in some cases still were “Upregulators.” Similarly, by 16S analysis, few Actinobacterial taxa changed in relative abundance in response to the treatments. This suggests that these populations were actively utilizing carbohydrates and not appreciably changing their population sizes over the timescale we measured. A recent SIP study on forest soils found that Actinobacteriota only accumulated ^13^C after 21 days (98). Since SIP requires replication to increase enrichment of DNA (99), the authors hypothesized that this could be due to slow growth. Our results support this hypothesis, and indicate that populations with minimal growth can still be active and functionally relevant in the community (100). We also note that some Actinobacterial populations in the Detritusphere guild had significant d-CAZy transcription as early as 3 days. Thus metatranscriptomics provides a way to assess functional relevance that is independent of changes in taxonomic relative abundance.

## CONCLUSIONS

Niche differentiation is central to theories of coexistence (25, 32, 83). Recent advances in metagenomic sequencing have allowed us to define the fundamental metabolic niches of representatives from poorly-known phyla, for whom there are little phenotypic data (57, 101). Using genome-centric metatranscriptomics to define the realized metabolic niches for soil populations, we found that carbohydrate depolymerization guilds rapidly emerged during rhizosphere community assembly. Using these guilds, we determined the prevalence of three d-CAZy transcriptional strategies in the rhizosphere, and found that rhizosphere organisms upregulate decomposition transcripts in addition to increasing population sizes. Further, these populations used both primary and breakdown products, and supports recent observations that metabolic handoffs link together interacting members of microbial communities (79–81). Guilds dynamics of carbohydrate depolymerization during rhizosphere succession provides a key step towards developing microbially-constrained models to predict the fate of soil carbon.

## Supporting information

Supplemental Methods and Results

## ACKNOWLEDGEMENTS

This research was supported by the U.S. Department of Energy Office of Science, Office of Biological and Environmental Research Genomic Science program under Awards SCW1589 and SCW1039 to JPR, DE-SC0010570 and DOE-SC0016247 to MKF, and DOE-SC10010566 to JFB. Sequencing was conducted as part of Community Sequencing Awards 1487 to JPR and 1472 to MKF. Work conducted at Lawrence Livermore National Laboratory was supported under the auspices of the U.S. Department of Energy under Contract DE-AC52-07NA27344. Work conducted at Lawrence Berkeley National Laboratory and the U.S. DOE. Joint Genome Institute, a DOE Office of Science User Facility, was supported under Contract No. DE-AC02-05CH11231. Cell imaging was conducted at the CNR Biological Imaging Facility at UC Berkeley. We thank Shengjing Shi, Katerina Estera, Jia Tian for assisting with sample processing and Alex Probst for providing bioinformatics support and advice.

