## Supplemental Methods and Results for "Niche differentiation is spatially and temporally regulated in the rhizosphere"

### SUPPLEMENTAL INFORMATION

### METHODS

#### Metatranscriptomic Sequencing (RNASeq)

Metatranscriptome libraries were prepared and sequenced at the Joint Genome Institute. Ribosomal RNA was depleted from 1 µg of total RNA using the Ribo-Zero rRNA Removal Kit (Epicentre) for Plants and Bacteria. Stranded cDNA libraries were generated using the Illumina TruSeq Stranded RNA LT kit. The rRNA depleted RNA was fragmented and reversed transcribed using random hexamers and SSII (Invitrogen) followed by second strand synthesis. The fragmented cDNA was treated with end-pair, A-tailing, adapter ligation, and 10 cycles of PCR. qPCR was used to determine the concentration of the libraries. Libraries were sequenced on the Illumina Hiseq.

The prepared libraries were quantified using KAPA Biosystem's next-generation sequencing library qPCR kit and run on a Roche LightCycler 480 real-time PCR instrument. The quantified libraries were then multiplexed into pools of 1-3 libraries each, and the pool was then prepared for sequencing on the Illumina HiSeq sequencing platform utilizing a TruSeq paired-end cluster kit, v3, and Illumina's cBot instrument to generate a clustered flowcell for sequencing. Sequencing of the flowcell was performed on the Illumina HiSeq2000 sequencer using a TruSeq SBS sequencing kit, v3, following a 2x150 indexed run recipe.

#### Single Amplified Genome (SAG) Sequencing

Single amplified genomes (SAGs) were generated following standard procedures in the Department of Energy Joint Genome Institute workflow (Rinke et al. 2014). Individual cells were sorted on a BD Influx (BD Biosciences) and treated with Ready-Lyse lysozyme (Epicentre; 5 U/µl final concentration) for 15 min at room temperature prior to the addition of lysis solution. Whole-genome amplification was performed with the REPLI-g Single Cell Kit (Qiagen) in 2 µl reactions set up with an Echo acoustic liquid handler (Labcyte). Amplification reactions were terminated after 6 h. Sequencing libraries were generated using the Nextera XT v2 kit (Illumina), and 2X150bp sequencing reads were sequenced on the Illumina Nextseq platform.

**Amplicon Sequencing (16S and ITS iTags)**

Each RNA sample prepared for metatranscriptomics sequencing was also reverse transcribed to create a paired cDNA sample for 16S and ITS amplicon analysis. 400 ng of TURBO DNase-treated RNA was reverse transcribed using the SuperScript III First-Strand Synthesis System (Invitrogen) with random hexamers according to the manufacturer's protocol. Plate-based 16S V4 region and ITS iTag preps are performed on the PerkinElmer Sciclone NGS robotic liquid handling system using 30 ng sample input, custom designed target primers with incorporated Illumina sequencing adapters. PCR amplicons were created using 25ul reactions using 10 ng cDNA, 1x 5PRIME HotMasterMix (Quantabio), 0.4 mg/ml BSA, and 200 nmol each primers (515F GTGCCAGCMGCCGCGGTAA; 805R GGACTACHVGGGTWTCTAAT) or Fungal ITS2 primers (ITS9 GAACGCAGCRAAIIGYGA; ITS4 TCCTCCGCTTATTGATATGC). Reverse primers were barcoded. Amplification procedure was as follows: initial denaturing at 94°C (3 min), 30 cycles of 94°C (45 sec), 50°C (60 sec), 72°C (90 sec), with a final extension at 72°C (10 min). After library sample prep, the samples from each target primer set were pooled together and the pool quantified using KAPA Biosystem's next-generation sequencing library qPCR kit and run on a Roche LightCycler 480 real-time PCR instrument. The pool was then loaded and sequenced on the Illumina MiSeq sequencing platform utilizing a MiSeq Reagent Kit, v3 600 cycle, following a 2x300 indexed run recipe.

**FIGURES**

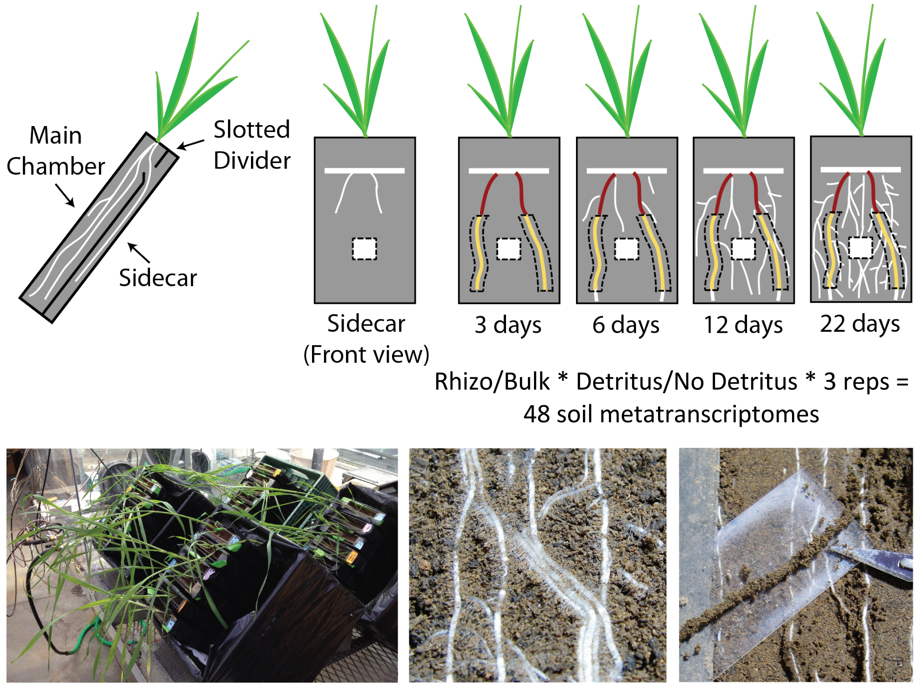


**Figure S1:** Experimental design. Plants were grown in microcosms with a separate experimental root chamber (sidecar). Root growth was tracked through the clear sidecar wall. After three days of root growth, individual root sections for all microcosms were marked to synchronize root age among all the microcosms. Marked root sections were destructively sampled at 3, 6, 12, and 22 days.

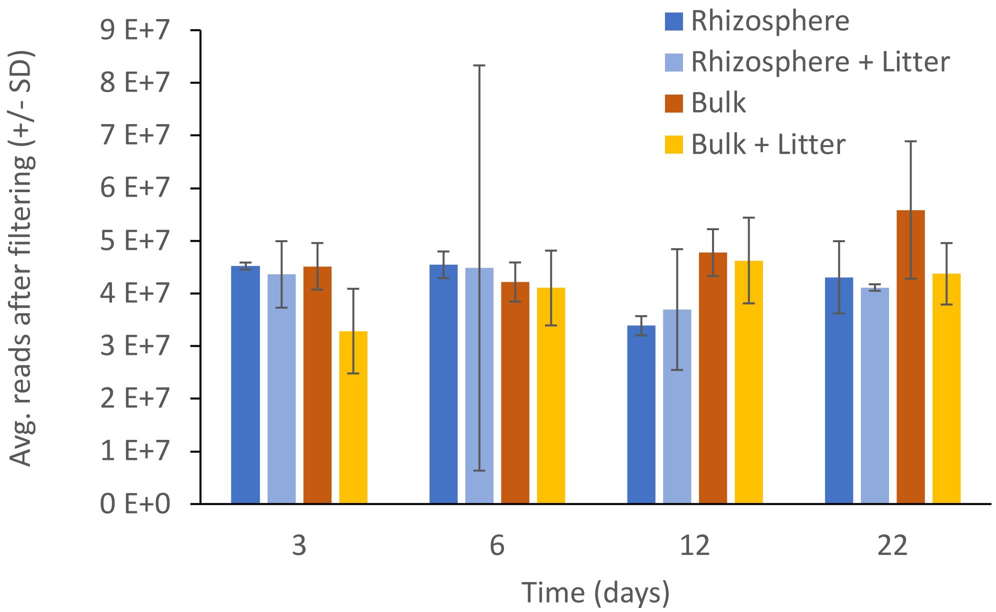


**Figure S2:** Average number of potential mRNA reads per sample after removing sequencing contaminants and ribosomal RNA reads (+/- standard deviation). Reads are averaged by treatment over time (3, 6, 12, 22 days) after exposure to the root (Rhizosphere, blue), bulk soil (Bulk, red), and amending both these habitats with root detritus (Rhizosphere + Detritus, light blue; Bulk + Detritus, yellow).

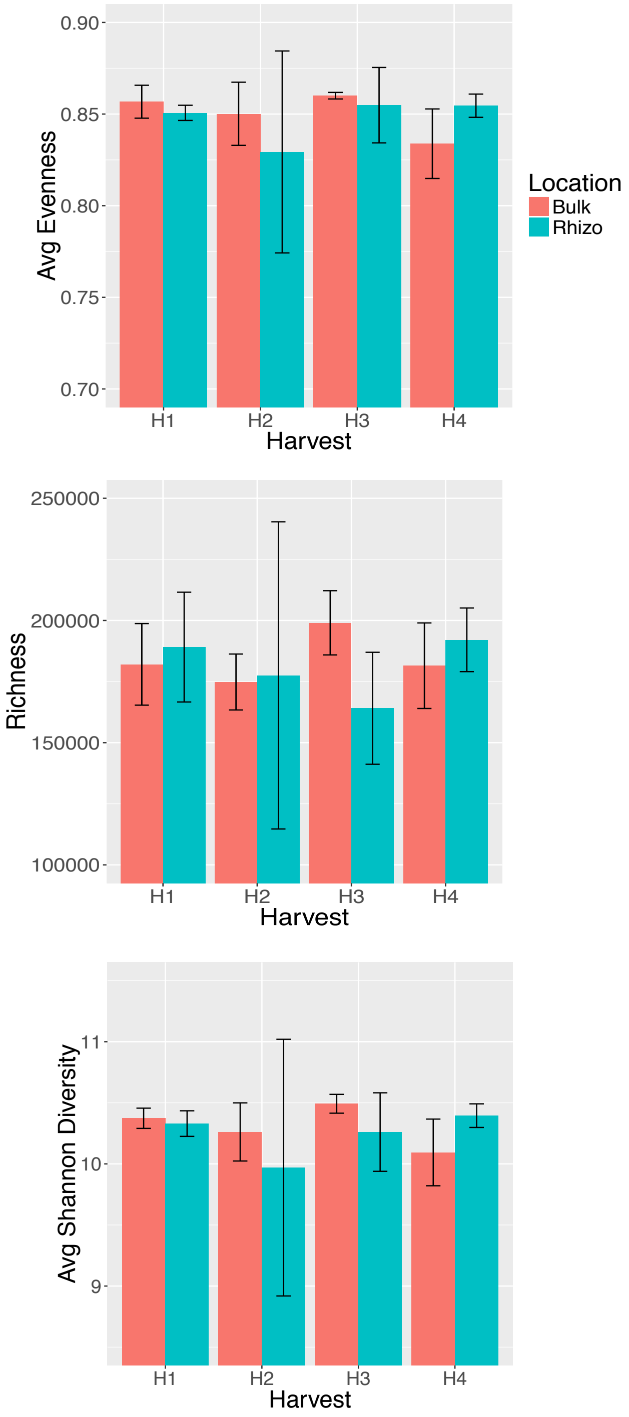


**Figure S3:** Average mRNA diversity after mapping to reference database for all rhizosphere (blue) and bulk soil (red) samples over time. Diversity is presented as average evenness (A), average richness (B), and average Shannon diversity (C) +/- standard deviation. H1-H4 refers to timepoints 3, 6, 12, 22 days, respectively.

A. B.

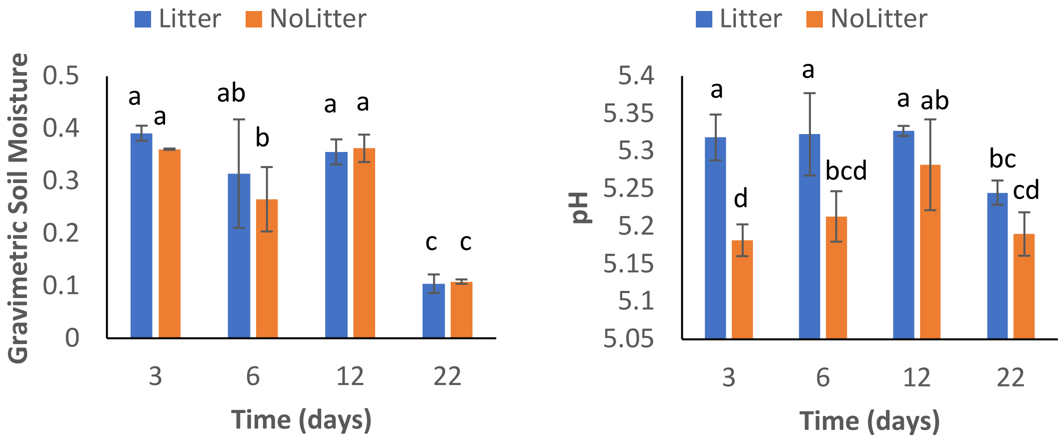


**Figure S4:** Average gravimetric soil moisture (A) and pH (B) of the microcosm soil amended with root detritus (blue) and without root detritus (orange) at each time point (+/- standard deviation). Letters represent Tukey HSD post hoc significance groups.

**A.** **B.**

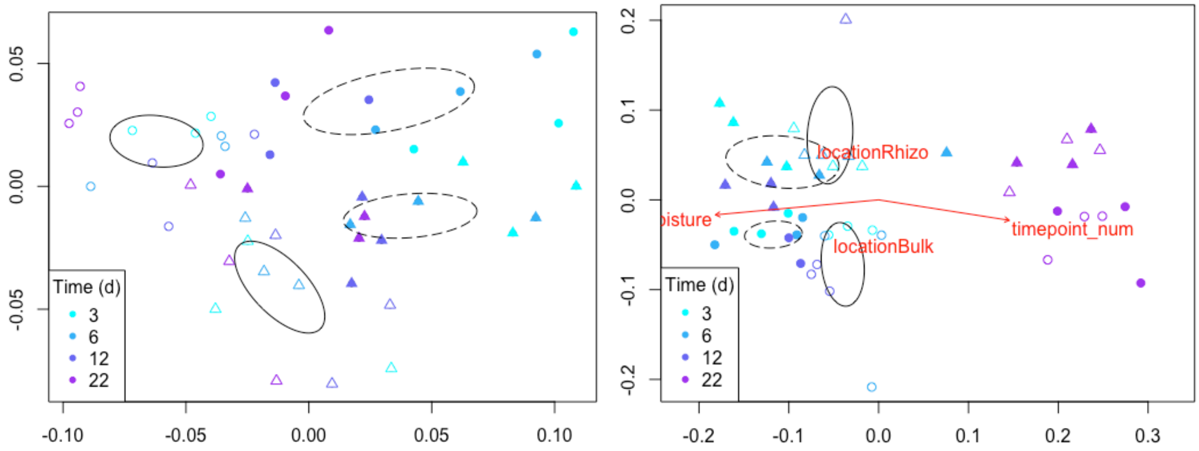


**Figure S5:** Influence of time 16S bacterial communities (A) and mRNA transcripts (B). Symbols represent the different treatments: Rhizosphere (hollow triangles), Rhizosphere + Detritus (filled triangles), Bulk (hollow circles), Bulk + Detritus (filled circles). Points are shaded by time point from light blue (3 days) to purple (22 days). Arrow overlays on indicate significant factors (Rhizosphere, Bulk) and numerical correlates (moisture, time) driving the community composition, as calculated by envfit.

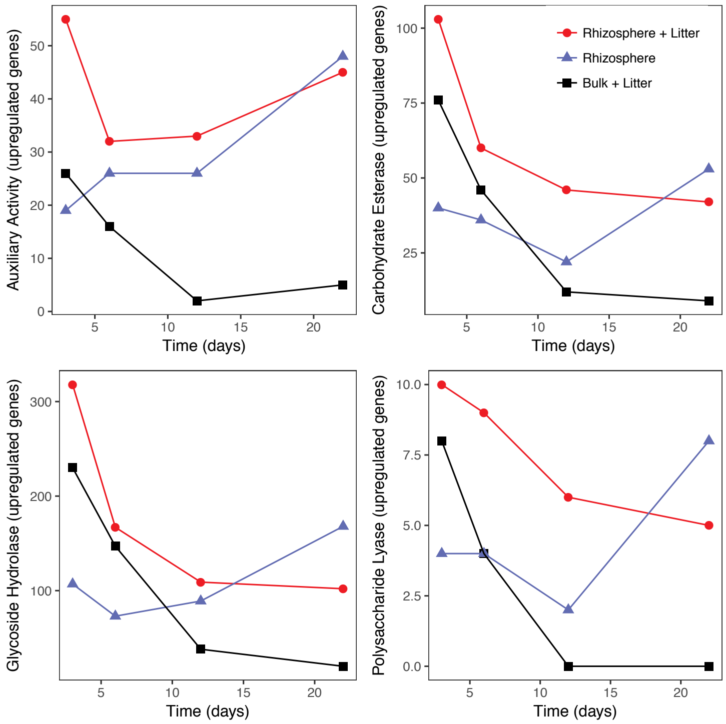


**Figure S6:** The cumulative number of significantly upregulated CAZy genes in the treatments relative to bulk soil for four CAZy enzyme classes (auxiliary activity, carbohydrate esterases, glycoside hydrolases, polysaccharide lyases) over the time-course experiment. The treatments are rhizosphere samples that contain detritus (red circles), bulk soil samples that contain detritus (black square), and rhizosphere without detritus addition (blue triangle).

**Supplemental Figure S7:** Population metatranscriptomes for carbohydrade degradation CAZymes for the 26 populations (A-Z below) that had 4 or more CAZymes significantly expressed. Gene annotation is presented in rows (CAZyme number and consensus annotation). Treatments are Rhizosphere (R), Rhizosphere + Detritus (RL), and Bulk Soil + Detritus (BL). H1-H4 represents the timepoints (3, 6, 12, 22 days, respectively). "S.P." indicates that the reference gene sequence had a potential signal peptide as calculated by SignalP (Petersen et al. 2011). 226 of 407 enzymes had recognizable signal peptides.

#### A. Burkholderiales 54_8 (Rhizosphere guild)

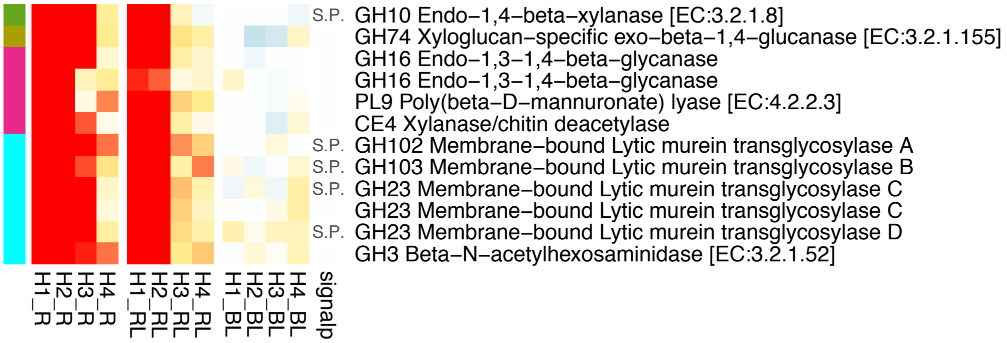


#### B. Janthinobacterium 65_15 (Rhizosphere guild)

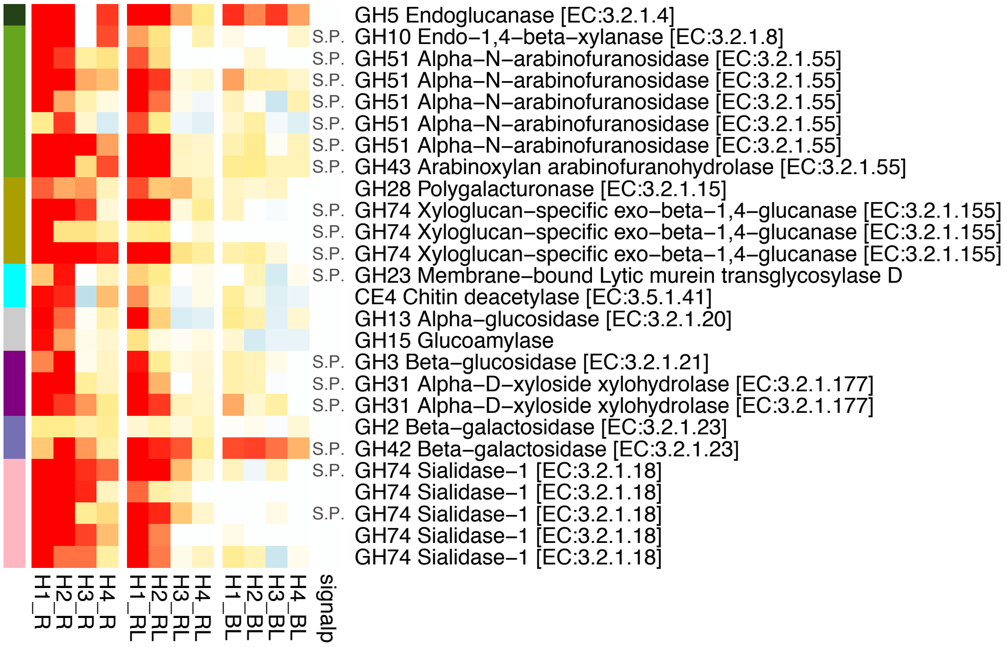


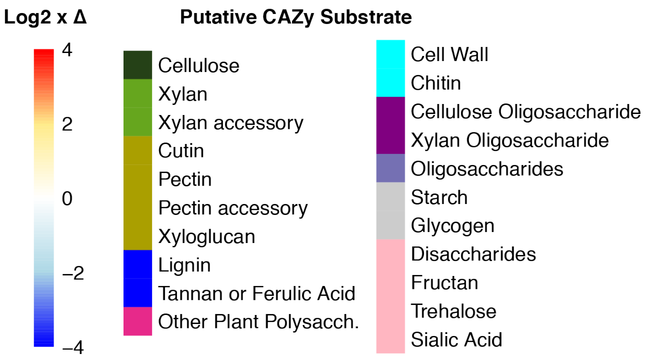

### C. Massilia 66_12 (Rhizosphere guild)

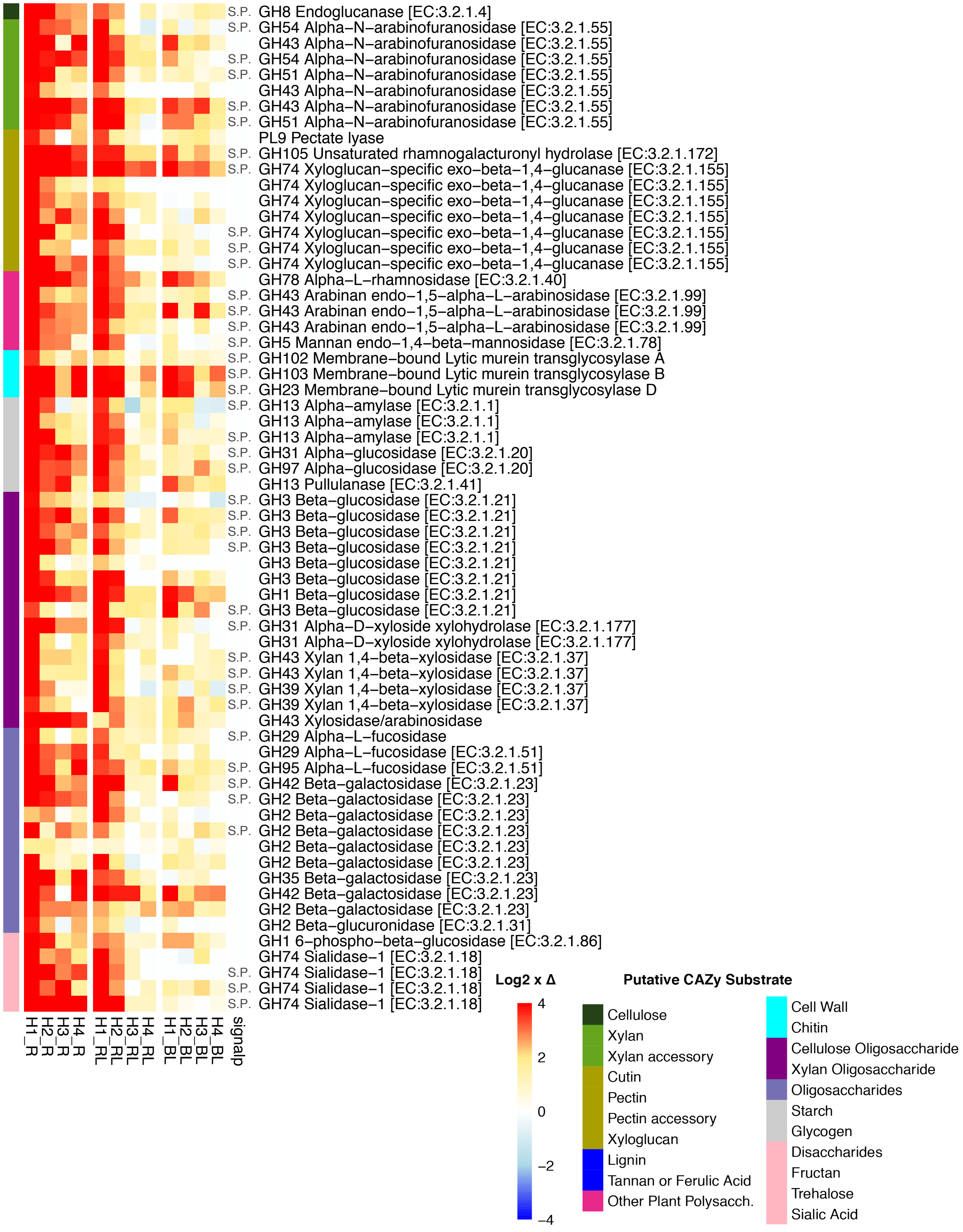

### D. Verrucomicrobiota 63_30 (Rhizosphere guild)

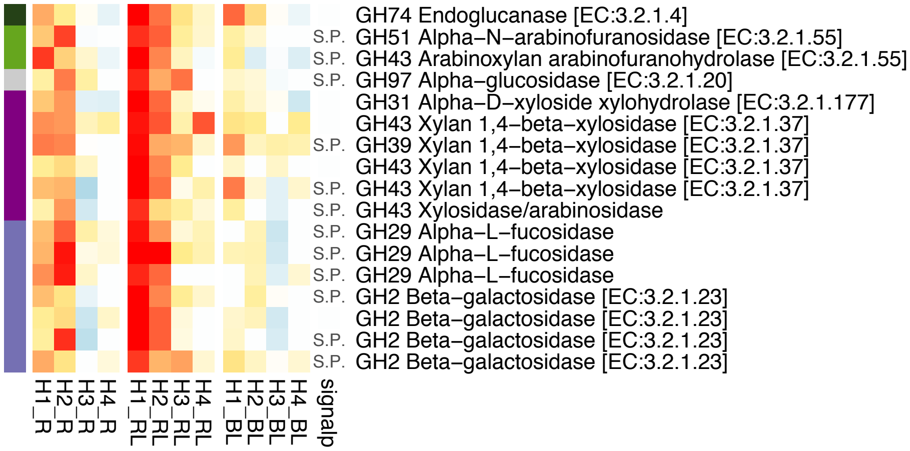


### E. Burkholderiales 62_29 (Rhizosphere guild)

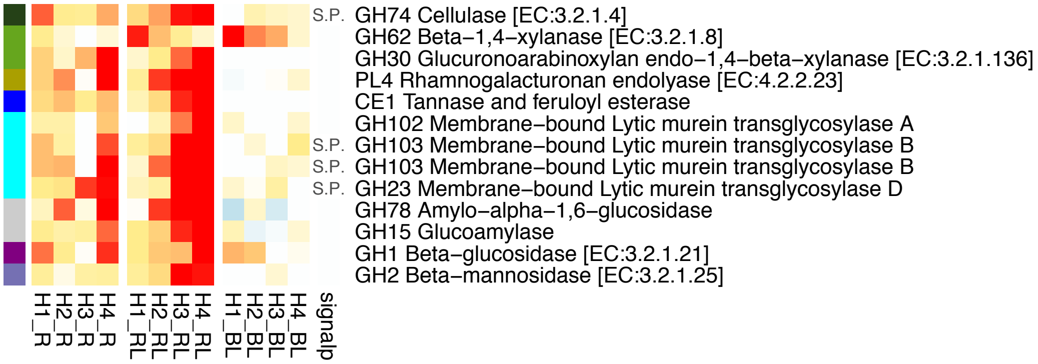


### F. Asticcacaulis 63_11 (Detritusphere guild)

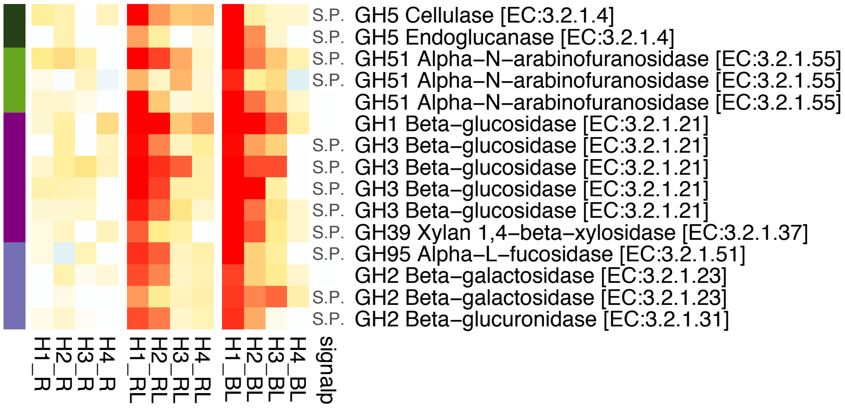


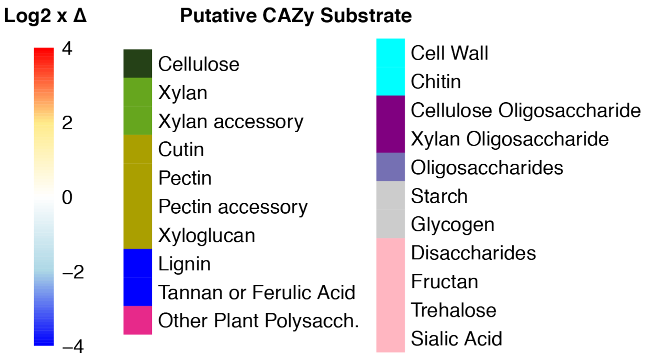

**G. Myxococcales 67_7 (Detritusphere guild)**

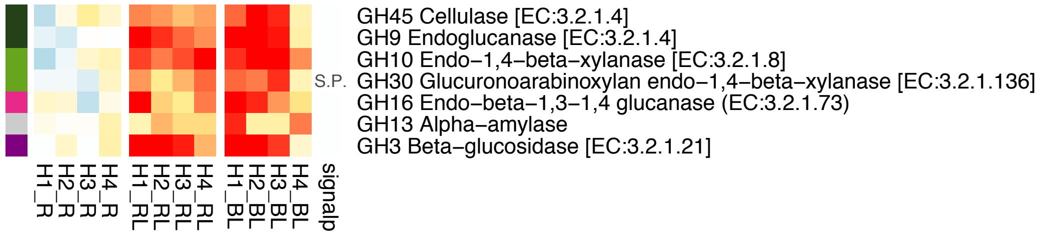


**H. Fibrobacterota 63_12 (Detritusphere guild)**

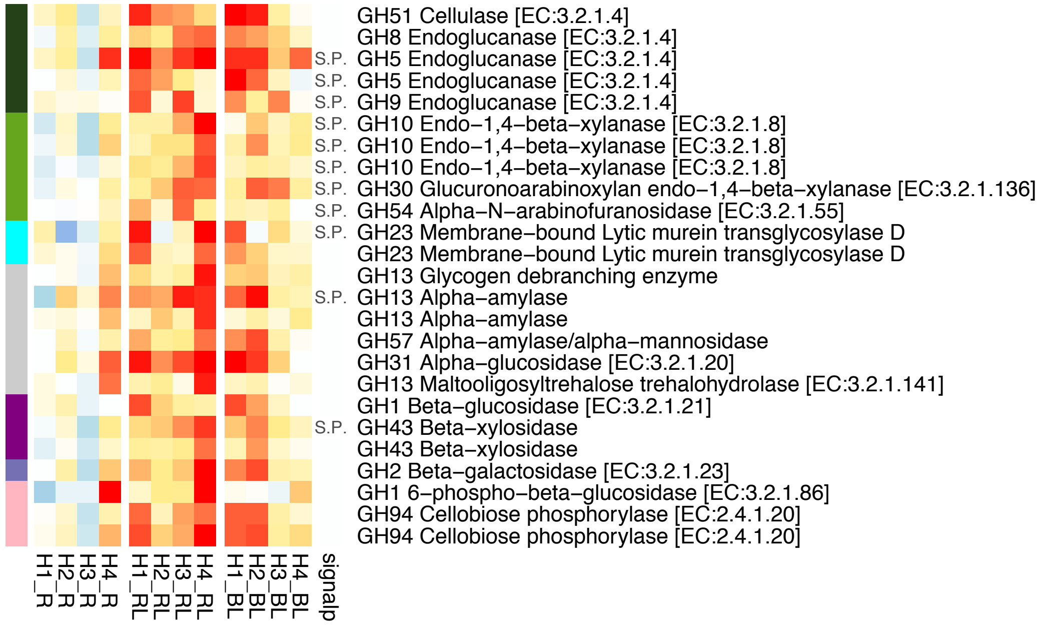


**I. Segetibacter b85 (Detritusphere guild)**

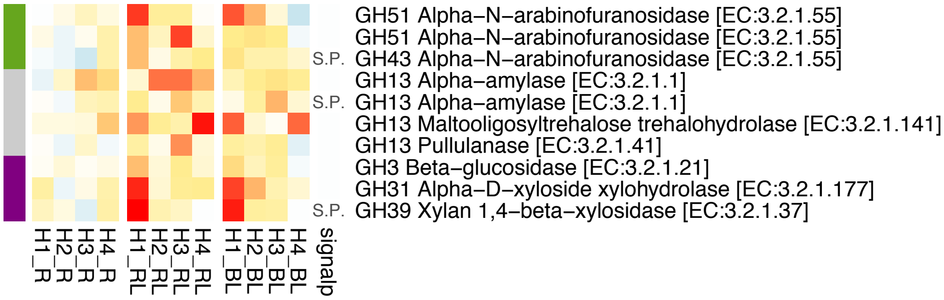


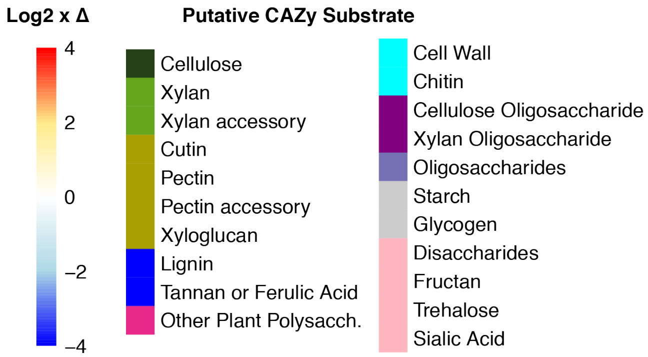

### J. Devosia JGI 030 (Detritusphere guild)

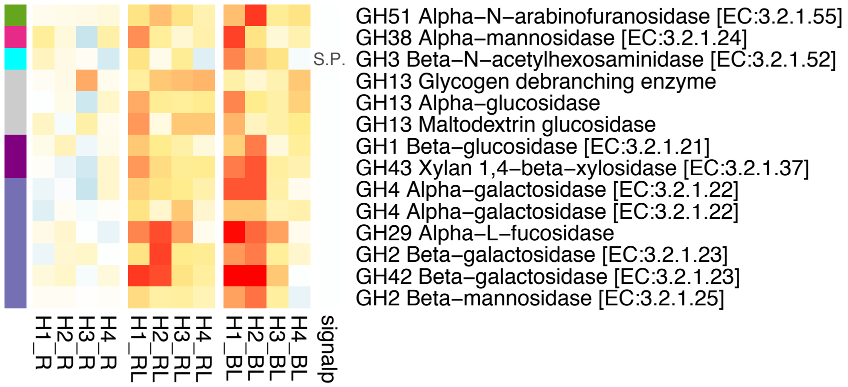


### K. Niastella b58 (Detritusphere guild)

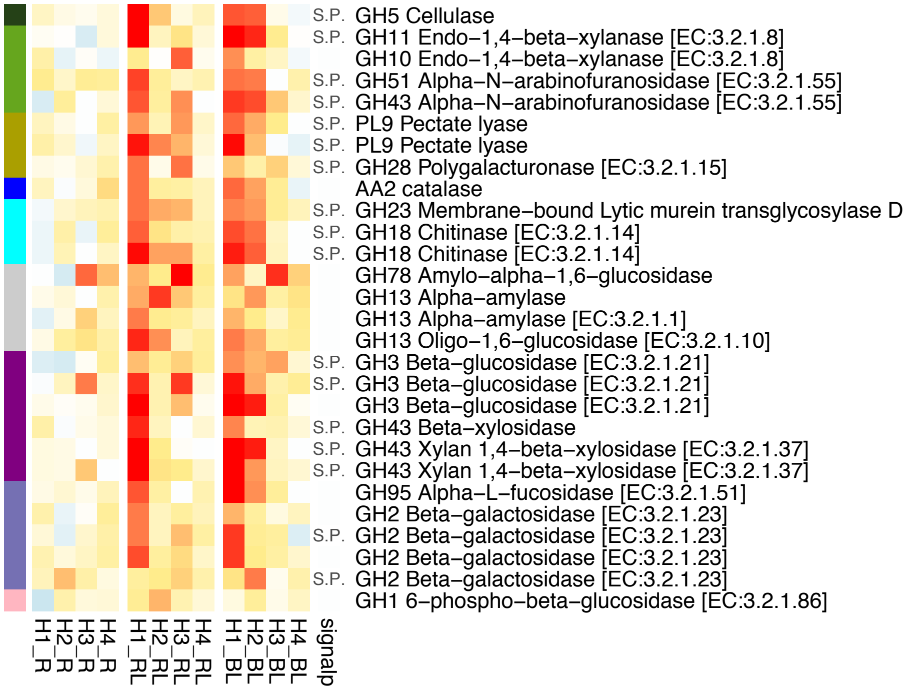


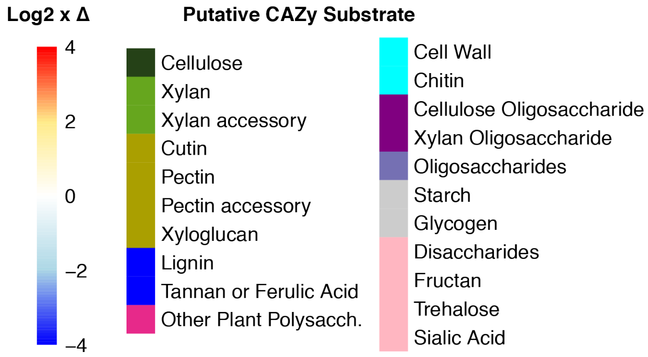

**L. Rhizobium 59_9 (Detritusphere guild)**

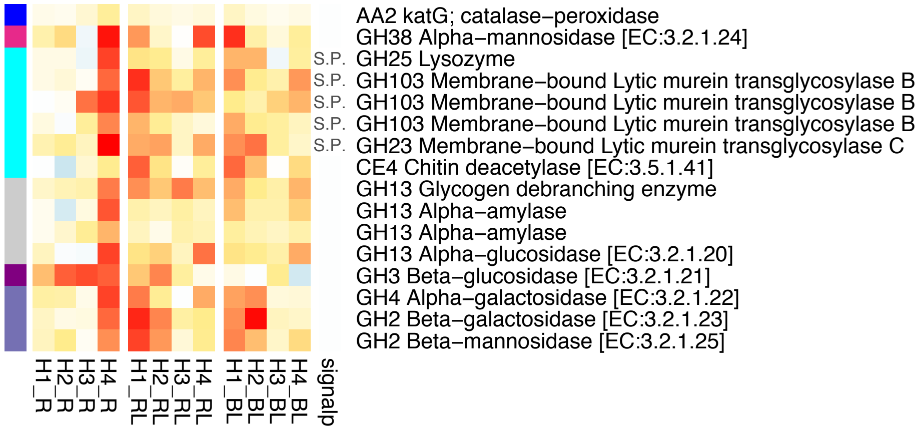


**M. Streptomyces tu. 70_12 (Detritusphere guild)**

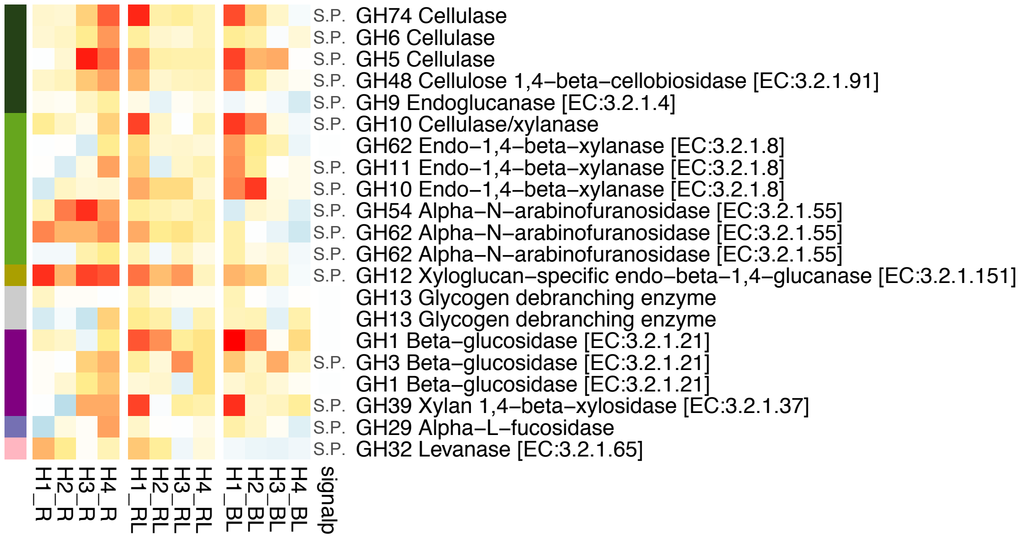


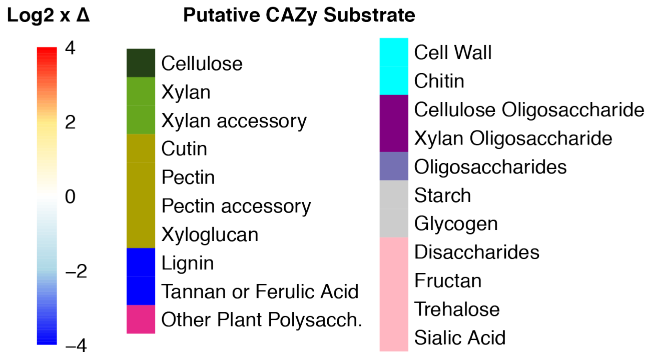

### N. Streptomycetales 68_8 (Detritusphere guild)

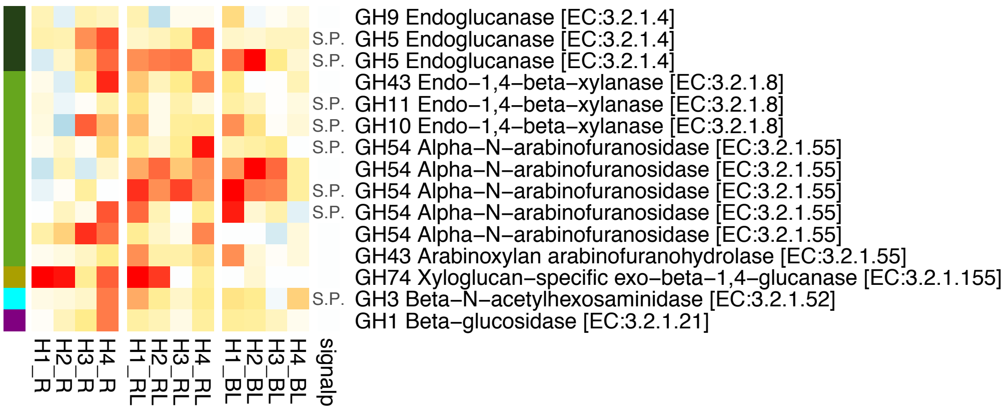


### O. Steptomyces 70_9 (Aging root guild)

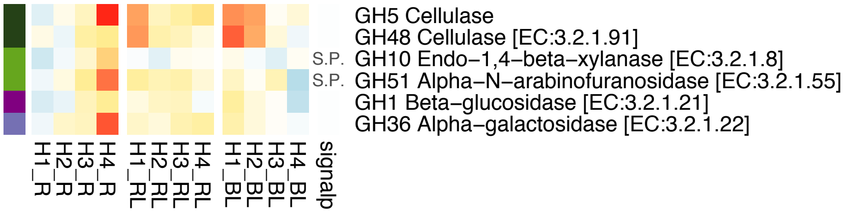


### P. Steptomyces 71_13 (Aging root guild)

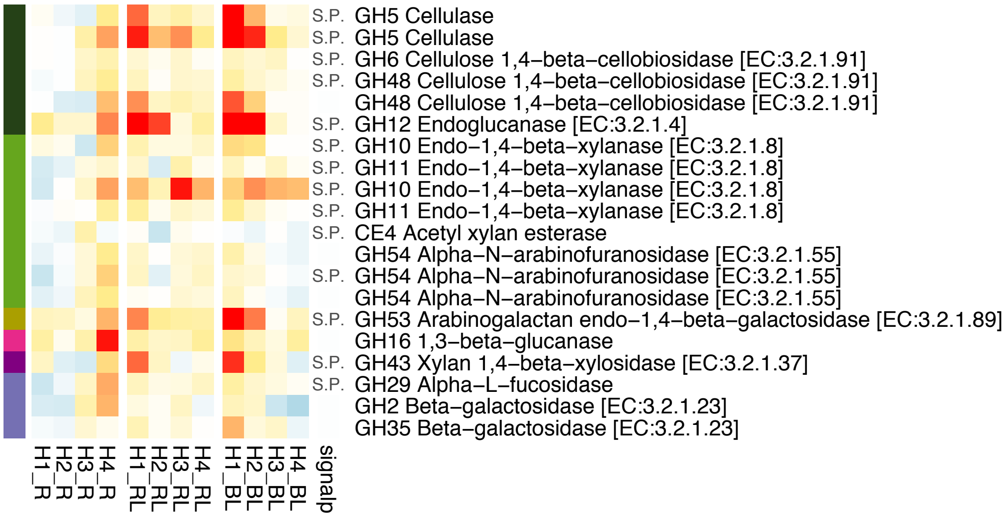


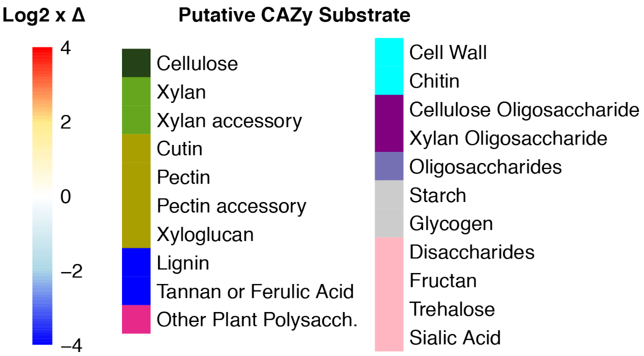

### Q. Streptomyces 71_16 (Aging root guild)

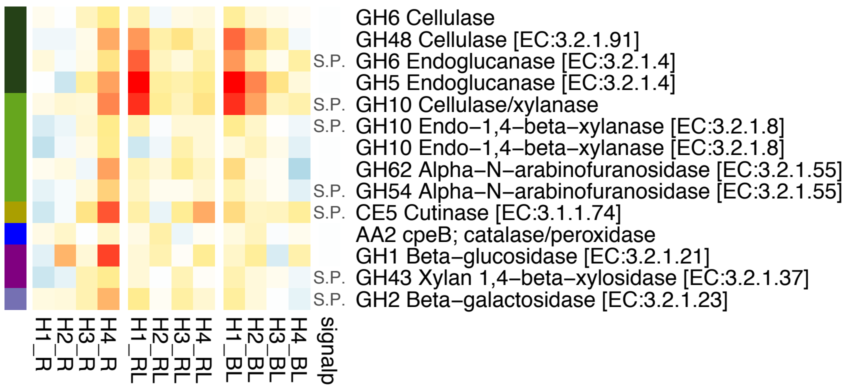


### R. Streptomyces 70_12 (Aging root guild)

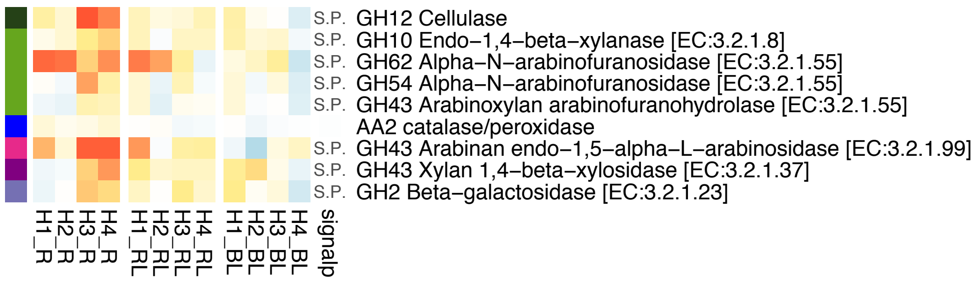


### S. Actinobacteriota 70_8 (Aging root guild)

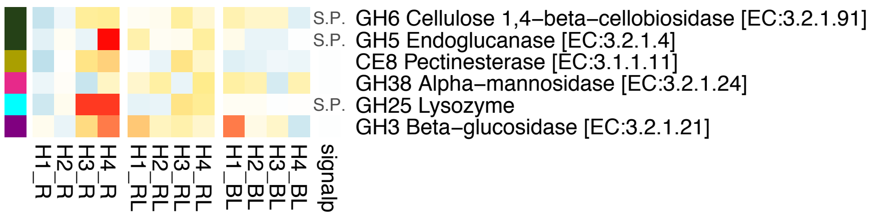


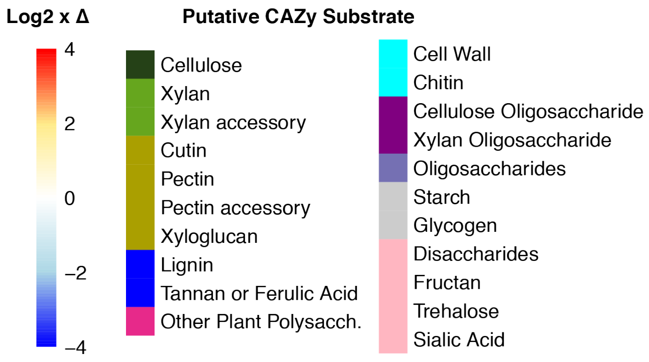

### T. Catenulisporales 70_18 (Aging root guild)

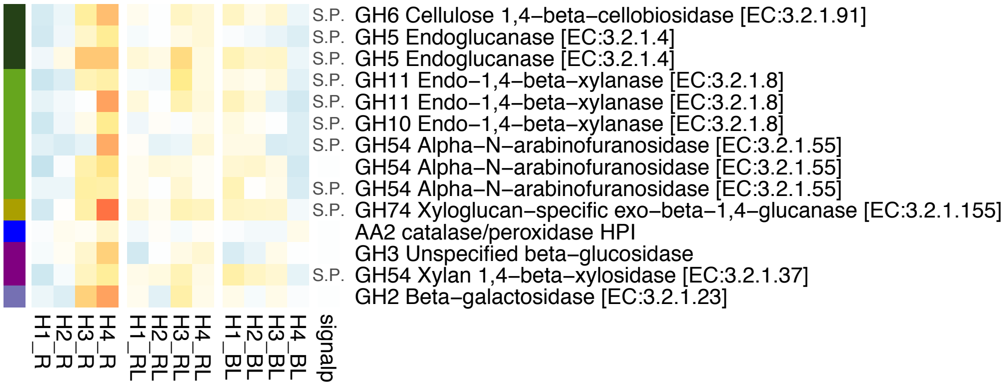


### U. Streptomyces URHA0041 (Aging root guild)

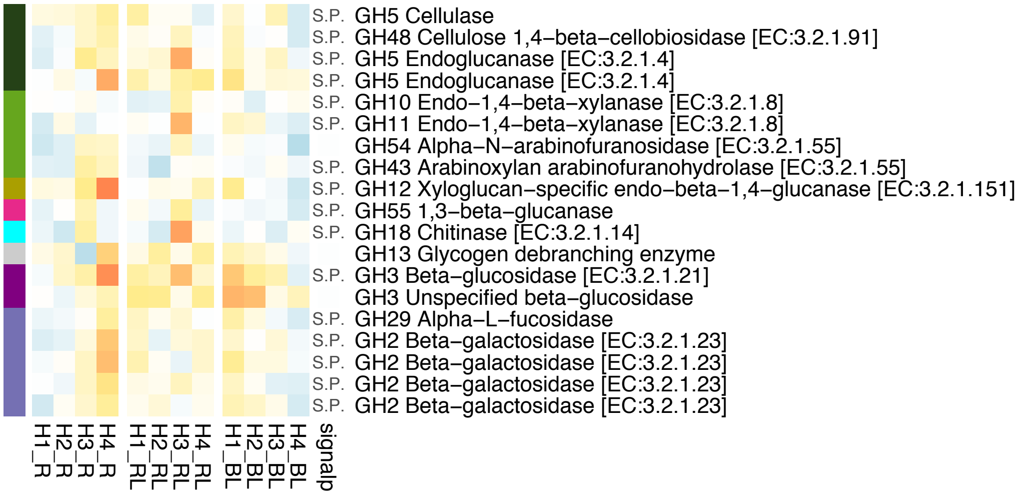


### V. Gammaproteobacteria b1 (Low response)

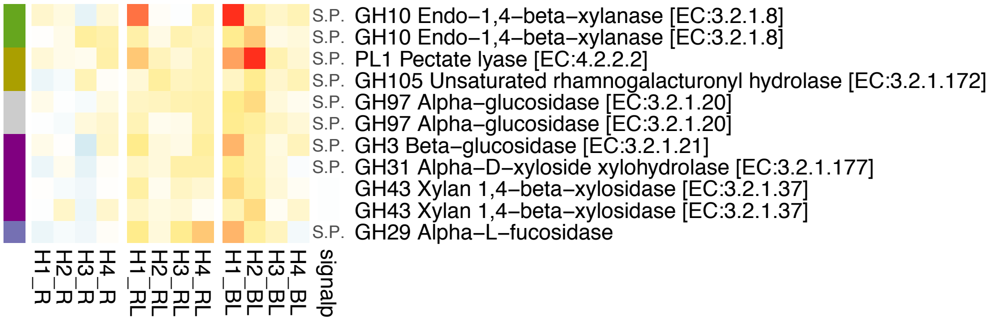


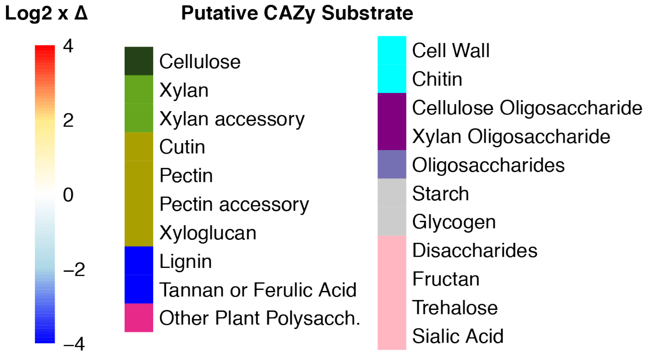

### W. Niabella b75 (Low response)

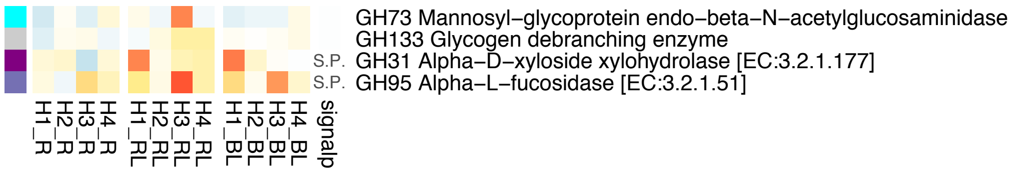


### X. Micromonosporaceae (Low response)

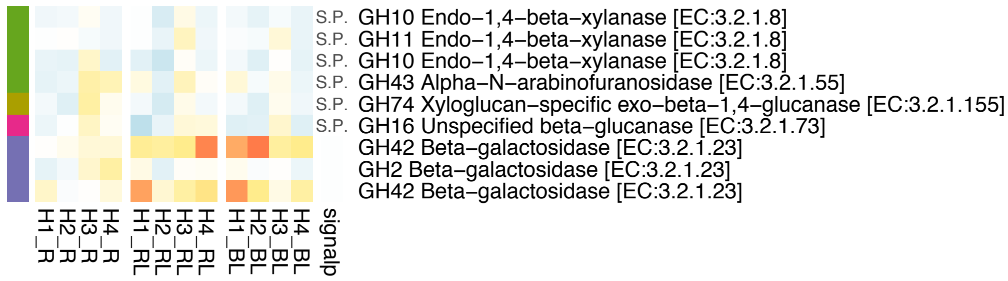


### Y. Bradyrhizobium URHA0013 (Low response)

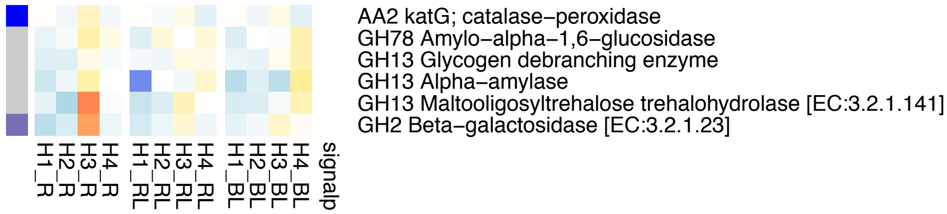


### Z. Kribbella b6 (Low response)

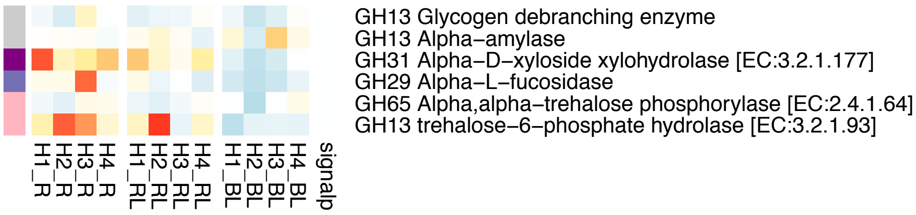


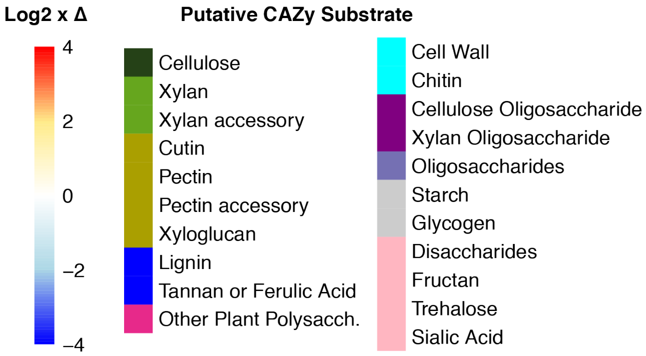

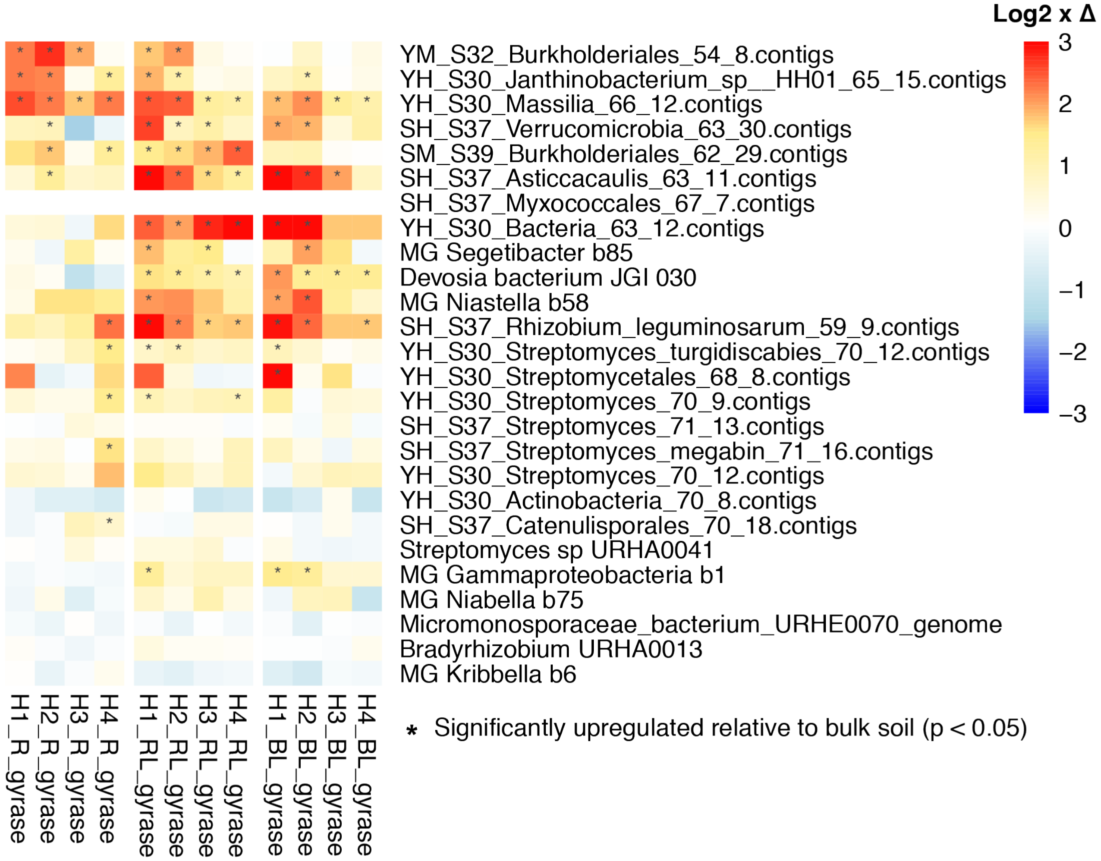


**Figure S8.** Population increases based on housekeeping gene expression (gyrase A, B), or "Growers." Heatmaps colors indicate the log2 fold increase (red) or decrease (blue) of average gyrase gene expression per genome relative to bulk soil over time (3, 6, 12, 22 days). Stars (*) indicate significant upregulation of gene expression relative to bulk soil by DESeq2. No gyrase data was available for Myxococcalis 67_7.

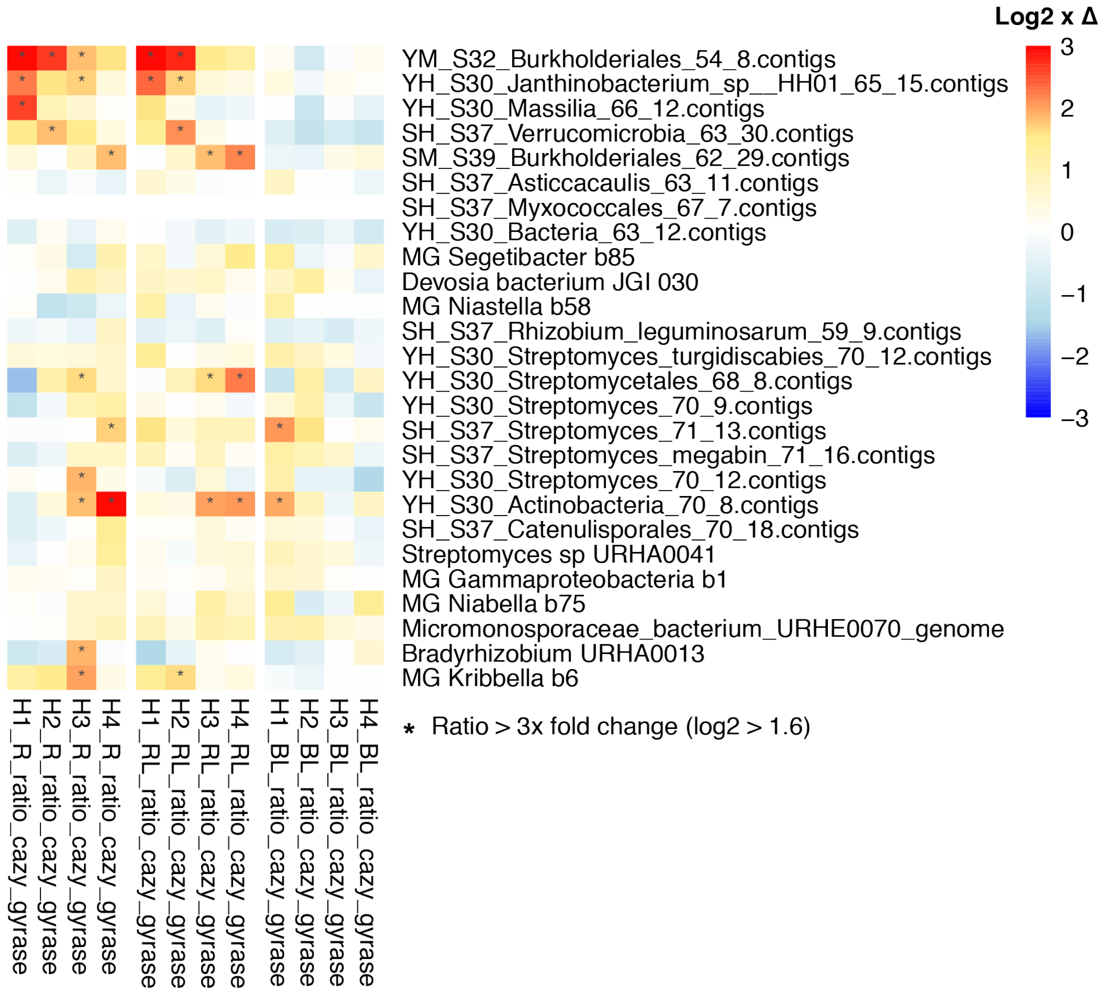


**Figure S9.** Population abundance-normalized CAZyme expression depicting "Upregulators." Heatmaps colors indicate the log2 fold increase (red) or decrease (blue) of average CAZyme gene expression per genome relative to housekeeping gene expression (gyrase A, B) over time (3, 6, 12, 22 days). Stars (*) indicate > 1.6x log2 fold change (i.e., 3-fold change). No gyrase data was available for Myxococcalis 67_7.

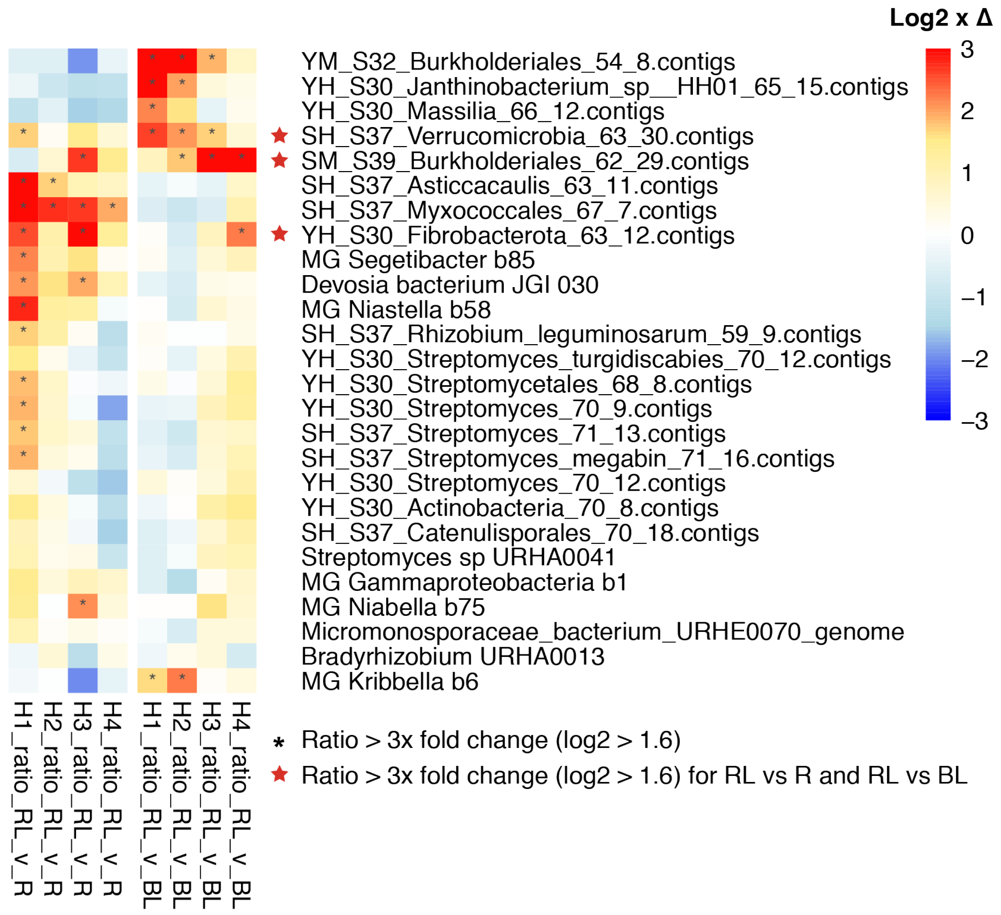


**Figure S10.** Identification of "Synergist" populations that have highest gene expression in the Rhizosphere + Detritus (RL) treatment relative to both the Rhizosphere (R) and Bulk + Detritus (BL) treatments. Heatmaps colors indicate the log2 fold increase (red) or decrease (blue) over time (3, 6, 12, 22 days). Stars (*) indicate > 1.6x log2 fold change (i.e., 3-fold change). Red stars indicate synergist populations that had a star (*) for both the RL vs R comparison and the RL vs BL comparison.

**Table S4:** F tables

**A.**

| 16S cDNA | F | $r^2$ | p | |
|---|---|---|---|---|
| Habitat | 16 | 0.15 | <0.0001 | *** |
| Detritus | 24 | 0.23 | <0.0001 | *** |
| Time | 6.5 | 0.19 | <0.0001 | *** |
| Detritus:Time | 2.2 | 0.063 | 0.021 | * |

**B.**

| ITS cDNA | F | $r^2$ | p | |
|---|---|---|---|---|
| Detritus | 19 | 0.25 | <0.0001 | *** |
| Time | 2.5 | 0.098 | <0.0001 | *** |

**C.**

| mRNA (3-22d) | F | $r^2$ | p | |
|---|---|---|---|---|
| Time | 3.2 | 0.16 | 0.0068 | ** |
| Habitat:Time | 3.0 | 0.052 | 0.011 | * |
| Habitat:Detritus:Time | 2.1 | 0.11 | 0.016 | * |

| mRNA (3-12d) | F | $r^2$ | p | |
|---|---|---|---|---|
| Habitat | 5.8 | 0.13 | <0.0001 | *** |
| Detritus | 5.1 | 0.11 | <0.0001 | *** |
| Time | 2.9 | 0.066 | <0.0001 | *** |
| Habitat:Time | 1.7 | 0.037 | 0.029 | * |
| Detritus:Time | 1.7 | 0.039 | 0.027 | * |

**Table S5:** Number of significantly upregulated bacterial OTUs relative to bulk soil by 16S itag sequence data. Data is aggregated at the family taxonomic level. Comparisons are between each treatment—Rhizosphere, Rhizosphere + Detritus, and Bulk Soil + Detritus—relative to Bulk soil without detritus amendment. Significance criteria are p<0.05 calculated by DESeq2 and accounts for multiple comparisons.

| | | Upregulated (Relative to Bulk) | | | | | | | | | | | |
|---|---|---|---|---|---|---|---|---|---|---|---|---|---|
| | | Rhizosphere | | | | Rhizosphere + Detritus | | | | Bulk Soil + Detritus | | | |
| **Phylum** | **Family** | 3 | 6 | 12 | 22 | 3 | 6 | 12 | 22 | 3 | 6 | 12 | 22 |
| **Acidobacteria** | Unknown | | | | | | | | | | | | |
| | Ellin6075 | | | | | | | | | | | | |
| | Koribacteraceae | | | | | | | | 1 | | | | |
| | RB40 | | | | | | | | 1 | | | | |
| | Solibacteraceae | | | | | | | | 1 | | | | |
| **Actinobacteria** | Unknown | | | | | | | | | | | | |
| | Actinosynnemataceae | | | | | | | | 1 | | | | |
| | AK1AB1_02E | | | | | | | | | | | 1 | |
| | Gaiellaceae | | | | | | | | | | | | |
| | Microbacteriaceae | | | | 1 | 1 | 1 | 1 | 1 | 1 | 1 | 3 | 1 |
| | Micrococcaceae | | | | | 1 | 1 | | | 1 | 1 | 1 | |
| | Micromonosporaceae | | | | | | | | | 1 | | | |
| | Nocardiaceae | | | | | | | | | | | 1 | |
| | Patulibacteraceae | | | | | | | | | | | | |
| | Solirubrobacteraceae | | | | | | | | | | | | |
| | Sporichthyaceae | | | | 1 | | | | 1 | | | | |
| | Streptomycetaceae | | | | 2 | | | | | | | | |
| | (blank) | | | | | | | | | | | | |
| **Armatimonadetes** | Unknown | | | 1 | 1 | 2 | 1 | | 4 | 2 | 2 | 1 | 1 |
| | Fimbriimonadaceae | | | | 1 | 6 | 8 | 1 | 8 | 6 | 8 | 2 | 5 |
| | Armatimonadaceae | | | | | 1 | 1 | 2 | 2 | 1 | 1 | | |
| **Bacteroidetes** | Unknown | | | | 3 | 2 | 2 | 5 | 17 | 4 | 4 | 2 | 6 |
| | Amoebophilaceae | | | | | | | 1 | 2 | | | | 2 |
| | Chitinophagaceae | | | 2 | 5 | 15 | 14 | 11 | 13 | 12 | 15 | 8 | 9 |
| | Cryomorphaceae | | 1 | 2 | 2 | 2 | 2 | 2 | 2 | 2 | 2 | 2 | 2 |
| | Cytophagaceae | | 1 | 2 | 5 | 14 | 8 | 7 | 16 | 15 | 15 | 6 | 10 |
| | Flavobacteriaceae | | | | | | | | 1 | 1 | | | |
| | Sphingobacteriaceae | | | | 3 | 3 | 3 | | 4 | 5 | 5 | | |
| | (blank) | | | | | 1 | 1 | 2 | 5 | | | 1 | 3 |
| **Chlamydiae** | Criblamydiaceae | | | | | | 1 | | | | 1 | | |
| | Parachlamydiaceae | | | | | | | 1 | 2 | 1 | 1 | | 2 |
| | (blank) | | | 1 | | 1 | 1 | 1 | 2 | 2 | 2 | | 1 |
| **Chlorobi** | Unknown | | | | | | | | 1 | | | | 2 |

| | | | | | | | | | | | | | |
|---|---|---|---|---|---|---|---|---|---|---|---|---|---|
| **Chloroflexi** | Unknown | | | | | | | | | | | | |
| | A4b | | | | | 4 | 1 | 2 | 3 | 2 | 2 | 3 | 3 |
| | oc28 | | | | | 1 | | | 3 | 1 | | | |
| | Thermogemmatisporaceae | | | | | | | | | | | | |
| | (blank) | | | | | | | 1 | | | | | |
| **Cyanobacteria** | Unknown | | | | | | 1 | | | | 1 | | 1 |
| **FBP** | Unknown | | 1 | | 3 | 1 | 3 | | 4 | 4 | 3 | | 3 |
| **Fibrobacteres** | Unknown | | | 1 | 1 | 3 | 2 | 4 | 4 | 3 | 3 | 4 | 1 |
| **Firmicutes** | Bacillaceae | | | | | | | | | | | | |
| | Clostridiaceae | | | | | | | | | | | 1 | |
| | Paenibacillaceae | 1 | 1 | | 1 | | | | | 1 | | 1 | |
| | Planococcaceae | | | | | | | | | | | | |
| | (blank) | | | | | | | | | | | | |
| **Planctomycetes** | Unknown | | | | | 1 | | 3 | 6 | 1 | | | 2 |
| | Gemmataceae | | | | | | | | | | | | |
| | Pirellulaceae | | | | | | 1 | | | | 1 | | |
| **Proteobacteria** | Unknown | | | 3 | 2 | 7 | 6 | 11 | 16 | 9 | 8 | 5 | 15 |
| | Acetobacteraceae | | | | | | | | | | | | |
| | Alteromonadaceae | 1 | | 1 | 1 | 3 | 1 | 1 | 1 | 3 | 2 | 1 | 1 |
| | Bacteriovoracaceae | | | | 1 | 3 | 1 | | 1 | 3 | 1 | | 1 |
| | Bdellovibrionaceae | 1 | 1 | 2 | 1 | 2 | 1 | 3 | 3 | 2 | 3 | 1 | 1 |
| | Caulobacteraceae | | 1 | 4 | 6 | 8 | 7 | 7 | 7 | 7 | 8 | 7 | 6 |
| | Comamonadaceae | 2 | 3 | 5 | 5 | 7 | 6 | 6 | 7 | 2 | 2 | 1 | 1 |
| | Coxiellaceae | | | | | 2 | 1 | | 1 | 2 | 1 | 1 | 1 |
| | Geobacteraceae | | | | | 1 | | | | 2 | | 6 | |
| | Haliangiaceae | | | | | 1 | 1 | 3 | 3 | 1 | 1 | 1 | 2 |
| | Hyphomicrobiaceae | | | | | 1 | | 1 | 2 | 1 | 1 | 1 | |
| | Legionellaceae | | | | | | 2 | | 2 | | 3 | | 1 |
| | Methylocystaceae | | | | | | | | | | 1 | | |
| | Myxococcaceae | | | | | | | | | | | | |
| | OM27 | | | | | 2 | 1 | 1 | 2 | 1 | 2 | | 1 |
| | Oxalobacteraceae | 7 | 6 | 2 | 3 | 9 | 9 | 2 | 4 | 4 | 3 | 1 | 1 |
| | Phyllobacteriaceae | | | | 2 | 1 | 1 | 1 | 4 | 3 | 2 | | 1 |
| | Polyangiaceae | | | | | 1 | | 2 | 3 | 1 | 1 | 1 | 3 |
| | Pseudomonadaceae | | | | 1 | | | | | 1 | | 1 | |
| | Rhizobiaceae | | | | 3 | 4 | 3 | 3 | 4 | 4 | 4 | 3 | 3 |
| | Rhodocyclaceae | | | | 1 | | | | | | | | |
| | Rhodospirillaceae | | | | 3 | 3 | 2 | 1 | 5 | 3 | 2 | 2 | 3 |
| | Rickettsiaceae | | | | | | | 1 | 1 | | | | 2 |
| | Sinobacteraceae | | | 1 | 1 | 3 | 2 | 3 | 5 | 5 | 3 | 2 | 5 |

| | | | | | | | | | | | | | |
|---|---|---|---|---|---|---|---|---|---|---|---|---|---|
| | Sphingomonadaceae | | | 5 | 6 | 6 | 4 | 5 | 8 | 6 | 3 | 6 | |
| | Syntrophobacteraceae | | | | | | | | | | | | |
| | Xanthobacteraceae | | | | | 1 | 1 | | 1 | 1 | 1 | | 1 |
| | Xanthomonadaceae | | | | 1 | 3 | 1 | 2 | 2 | 8 | 3 | 1 | 2 |
| | (blank) | | 1 | 3 | 3 | 6 | 5 | 10 | 26 | 6 | 8 | 6 | 14 |
| **TM6** | Unknown | | | | | | 3 | | | 1 | 3 | | |
| **Verrucomicrobia** | Unknown | 1 | | 2 | 1 | 5 | 3 | 5 | 6 | 2 | 2 | 2 | 6 |
| | Chthoniobacteraceae | | 1 | 2 | 7 | 8 | 10 | 9 | 8 | 6 | 8 | 1 | 3 |
| | 01D2Z36 | | | | | 1 | 1 | 1 | 1 | 1 | 1 | | 1 |
| | Ellin515 | | | | | 2 | 1 | | 1 | 1 | 1 | | |
| | Ellin517 | | | | 3 | | | | 3 | | 1 | | |
| | Opitutaceae | 2 | 3 | 4 | | 3 | 3 | 5 | 5 | 2 | 1 | | 3 |
| | Verrucomicrobiaceae | | | | | 1 | | | | 1 | 2 | | |
| | (blank) | | | 1 | 1 | 3 | | | 3 | 2 | 1 | | 1 |
| **Unknown** | (blank) | | | | | 5 | 4 | 2 | 1 | 4 | 8 | 2 | 5 |
| **Total** | | 15 | 20 | 44 | 81 | 161 | 132 | 129 | 241 | 161 | 159 | 89 | 138 |

**Table S6:** Number of significantly downregulated bacterial OTUs relative to bulk soil by 16S itag sequence data. Data is aggregated at the family taxonomic level. Comparisons are between each treatment—Rhizosphere, Rhizosphere + Detritus, and Bulk Soil + Detritus—relative to Bulk soil without detritus amendment. Significance criteria are p<0.05 calculated by DESeq2 and accounts for multiple comparisons.

| | | Downregulated (Relative to Bulk) | | | | | | | | | | | |
|---|---|---|---|---|---|---|---|---|---|---|---|---|---|
| | | Rhizosphere | | | | Rhizosphere + Detritus | | | | Bulk Soil + Detritus | | | |
| **Phylum** | **Family** | **3** | **6** | **12** | **22** | **3** | **6** | **12** | **22** | **3** | **6** | **12** | **22** |
| **Acidobacteria** | Unknown | | | | | | | 3 | | | 1 | | |
| | Ellin6075 | | | | | | | 2 | 1 | | | | |
| | Koribacteraceae | | | | | | | | 1 | 1 | | | |
| | RB40 | | | | | | | | | | | | |
| | Solibacteraceae | | | | | | | | | | | | |
| **Actinobacteria** | Unknown | | | | | | 1 | | 1 | | 2 | 1 | |
| | Actinosynnemataceae | | | | | | | | | | | | |
| | AK1AB1_02E | | | | | | | 1 | 1 | | | | |
| | Gaiellaceae | | | | 1 | 1 | 1 | 3 | 2 | | 1 | | 5 |
| | Microbacteriaceae | | | | | | | 1 | | | | | |
| | Micrococcaceae | | | | | | | | | | | | |
| | Micromonosporaceae | | | | | | | | | | 1 | | |
| | Nocardiaceae | | | | | | | | | | | | |
| | Patulibacteraceae | | | | | | | | 1 | | | | 1 |
| | Solirubrobacteraceae | | | | | | | | 2 | | 3 | | 2 |
| | Sporichthyaceae | | | | | | | | | | | 1 | 1 |
| | Streptomycetaceae | | | | | | | | | | | | |
| | (blank) | | | | | 1 | | | 1 | | 1 | | 1 |
| **Armatimonadetes** | Unknown | | | | | | | | | | | | |
| | Fimbriimonadaceae | | | | | | | 1 | | | | | |
| | Armatimonadaceae | | | | | | | | | | | | |
| **Bacteroidetes** | Unknown | | | | | | | | | | | | |
| | Amoebophilaceae | | | | | | | | | | | | |
| | Chitinophagaceae | | | | | | | | | | | | |
| | Cryomorphaceae | | | | | | | | | | | | |
| | Cytophagaceae | | | | | 1 | | 1 | 2 | 1 | | | |
| | Flavobacteriaceae | | | | | | | | | | | | |
| | Sphingobacteriaceae | | | | | | | | | | | | |
| | (blank) | | | | | | | | | | | | |
| **Chlamydiae** | Criblamydiaceae | | | | | | | | | | | | |
| | Parachlamydiaceae | | | | | | | | | | | 1 | |
| | (blank) | | | | | | | | | | | | |

| | | | | | | | | | | | | | |
|---|---|---|---|---|---|---|---|---|---|---|---|---|---|
| **Chlorobi** | Unknown | | | | | | | | | | | | |
| **Chloroflexi** | Unknown | | | | | | | | 4 | | | | 6 |
| | A4b | | | | | | | | | | | | |
| | oc28 | | | | | | | | | | | | |
| | Thermogemmatisporaceae | | | | | | | 1 | | | | | |
| | (blank) | | | | | | | | | | | | |
| **Cyanobacteria** | Unknown | | | | 1 | | 1 | 1 | | | 1 | 1 | |
| **FBP** | Unknown | | | | | | | | | | | | |
| **Fibrobacteres** | Unknown | | | | | | | | | | | | |
| **Firmicutes** | Bacillaceae | | | | | 1 | | 1 | | 1 | | | |
| | Clostridiaceae | | | | | | | | | | | | |
| | Paenibacillaceae | | | | | 2 | | 2 | 1 | | | | |
| | Planococcaceae | | | | | 1 | | | 1 | | | | |
| | (blank) | | | | | 1 | | 2 | 1 | | | | |
| **Planctomycetes** | Unknown | | | | | | | | 2 | | | | 1 |
| | Gemmataceae | | | | | | | | 1 | | | | |
| | Pirellulaceae | | | | | | | | | | | | |
| **Proteobacteria** | Unknown | | | | 1 | 3 | 2 | 1 | 3 | 5 | 2 | 1 | 1 |
| | Acetobacteraceae | | | | | | | 1 | | | | | |
| | Alteromonadaceae | | | | | | | | | | | | |
| | Bacteriovoracaceae | | | | | | | | | | | | |
| | Bdellovibrionaceae | | | | | | | | | | | | |
| | Caulobacteraceae | | | | | | | 1 | | | | | |
| | Comamonadaceae | | | | | | | | | | | | |
| | Coxiellaceae | | | | | | | | | | | | |
| | Geobacteraceae | | | | | | | | | | | | |
| | Haliangiaceae | | | | | | | | | | | | |
| | Hyphomicrobiaceae | | | | | | | | 1 | 1 | | | |
| | Legionellaceae | | | | | | | | | | | | |
| | Methylocystaceae | | | | | | | | | | | | |
| | Myxococcaceae | | | | | | 1 | | | | | | |
| | OM27 | | | | | | | | | | | | |
| | Oxalobacteraceae | | | | | | | | | | | | |
| | Phyllobacteriaceae | | | | | | | | | | | | |
| | Polyangiaceae | | | | | | | | | | | | |
| | Pseudomonadaceae | | | | | | | | | | | | |
| | Rhizobiaceae | | | | | | | | | | | | |
| | Rhodocyclaceae | | | | | | | | | | | | |
| | Rhodospirillaceae | | | | 1 | | | | | | | | |
| | Rickettsiaceae | | | | | | | | | | | 1 | |

| | | | | | | | | | | | | | |
|---|---|---|---|---|---|---|---|---|---|---|---|---|---|
| | Sinobacteraceae | | | | | | | 1 | | | | | |
| | Sphingomonadaceae | | | | | | | | | | | | |
| | Syntrophobacteraceae | | | | | | | | | 1 | | | |
| | Xanthobacteraceae | | | | | | | | | | | | |
| | Xanthomonadaceae | | | | | | | | | | | | |
| | (blank) | | | | | 1 | | | 4 | 1 | 1 | 1 | 1 |
| **TM6** | Unknown | | | | | | | | | | | | |
| **Verrucomicrobia** | Unknown | | | | | | | | 1 | | | | 1 |
| | Chthoniobacteraceae | | | | | | | 2 | 1 | | | 1 | |
| | 01D2Z36 | | | | | | | | | | | | |
| | Ellin515 | | | | | | | | | | | | |
| | Ellin517 | | | | | | | | | | | | |
| | Opitutaceae | | | | | | | | | | | | |
| | Verrucomicrobiaceae | | | | | | | | | | | | |
| | (blank) | | | | | | | | | | | | |
| **Unknown** | (blank) | | | | | | | 3 | 1 | | | 1 | |
| **Total** | | 0 | 0 | 0 | 4 | 12 | 6 | 28 | 33 | 11 | 13 | 9 | 20 |

**Table S7:** Decomposition CAZymes and their putative substrates used to assess the carbohydrate degradation potential for the population metatranscriptomes. The number of unique genes significantly up- or down-regulated relative to bulk soil was totaled per annotation category (all treatments aggregated). Significance was determined by DESeq2. Gene annotation is a consensus annotation from KEGG, ggkbase, and dbCAN (assigned CAZy numbers noted). References provided where relevant for clarification.

| Putative Substrate | Gene Annotation (Reference) | Assigned CAZy | # Sig. Genes |
|---|---|---|---|
| **Cell Wall** | Lysozyme | GH25 | 3 |
| | Membrane-bound Lytic murein transglycosylase A | GH102 | 3 |
| | Membrane-bound Lytic murein transglycosylase B | GH103 | 7 |
| | | GH23 | 1 |
| | Membrane-bound Lytic murein transglycosylase C | GH23 | 7 |
| | Membrane-bound Lytic murein transglycosylase D | GH23 | 12 |
| **Cellulose** | Cellulase | GH12 | 1 |
| | | GH5 | 7 |
| | | GH6 | 2 |
| | | GH74 | 1 |
| | Cellulase [EC:3.2.1.4] | GH45 | 1 |
| | | GH5 | 2 |
| | | GH51 | 1 |
| | | GH6 | 1 |
| | | GH74 | 1 |
| | Cellulase [EC:3.2.1.91] | GH48 | 2 |
| | Cellulose 1,4-beta-cellobiosidase [EC:3.2.1.91] | GH48 | 4 |
| | | GH6 | 3 |
| | Endoglucanase [EC:3.2.1.4] (Berlemont and Martiny 2013) | GH12 | 1 |
| | | GH5 | 13 |
| | | GH6 | 1 |
| | | GH74 | 1 |
| | | GH8 | 2 |
| | | GH9 | 4 |
| **Chitin** | Beta-N-acetylhexosaminidase [EC:3.2.1.52] (Konno et al. 2012) | GH3 | 4 |
| | Chitin deacetylase [EC:3.5.1.41] | CE4 | 2 |
| | Chitinase [EC:3.2.1.14] | GH18 | 4 |
| | Mannosyl-glycoprotein endo-beta-N-acetylglucosaminidase | GH73 | 1 |
| **Cutin** | Cutinase [EC:3.1.1.74] | CE5 | 1 |
| **Disaccharides** | 6-phospho-beta-glucosidase [EC:3.2.1.86] (Yu et al. 2013) | GH1 | 4 |
| | | GH4 | 1 |
| | Cellobiose phosphorylase [EC:2.4.1.20] | GH94 | 4 |
| **Fructan** | Levanase [EC:3.2.1.65] | GH32 | 1 |

| | | | |
|---|---|---|---|
| **Glycogen** | Amylo-alpha-1,6-glucosidase | GH78 | 4 |
| | Glycogen debranching enzyme | GH13 | 25 |
| | | GH133 | 2 |
| **Lignin** | catalase | AA2 | 2 |
| | catalase/hydroperoxidase HPI(I) | AA2 | 2 |
| | catalase/peroxidase | AA2 | 1 |
| | catalase/peroxidase HPI | AA2 | 3 |
| | catalase/peroxidase HPI (EC:1.11.1.6) | AA2 | 3 |
| | cpeB; catalase/peroxidase | AA2 | 1 |
| | katG; catalase (EC:1.11.1.6 1.11.1.7) | AA2 | 1 |
| | katG; catalase (EC:1.11.1.6) | AA2 | 1 |
| | katG; catalase peroxidase (EC:1.11.1.21) | AA2 | 1 |
| | katG; catalase-peroxidase | AA2 | 2 |
| | katG; Catalase-peroxidase (EC:1.11.1.21) | AA2 | 1 |
| | katG; catalase-peroxidase (EC:1.11.1.6) | AA2 | 1 |
| | katG; peroxidase/catalase katG (EC:1.11.1.7 1.11.1.8) | AA2 | 1 |
| | katGa; peroxidase/catalase oxidoreductase (EC:1.11.1.6) | AA2 | 1 |
| **Cellulose-Oligosaccharide** | Beta-glucosidase [EC:3.2.1.21] (Berlemont and Martiny 2013, Bergmann et al. 2014) | GH1 | 15 |
| | | GH3 | 31 |
| | Unspecified beta-glucosidase | GH3 | 2 |
| **Xylan-Oligosaccharide** | Alpha-D-xyloside xylohydrolase [EC:3.2.1.177] | GH31 | 10 |
| | Beta-xylosidase (Bajpai 1997) | GH39 | 2 |
| | | GH43 | 4 |
| | Oligosaccharide reducing-end xylanase [EC:3.2.1.156] | GH8 | 1 |
| | Xylan 1,4-beta-xylosidase [EC:3.2.1.37] | GH39 | 6 |
| | | GH43 | 15 |
| | | GH54 | 1 |
| | Xylosidase/arabinosidase | GH43 | 2 |
| **Oligosaccharides** | Alpha-galactosidase [EC:3.2.1.22] | GH36 | 1 |
| | | GH4 | 7 |
| | Alpha-L-fucosidase (Cao et al. 2014) | GH29 | 10 |
| | Alpha-L-fucosidase [EC:3.2.1.51] | GH29 | 1 |
| | | GH95 | 5 |
| | Beta-galactosidase [EC:3.2.1.23] | GH2 | 32 |
| | | GH35 | 2 |
| | | GH42 | 11 |
| | Beta-glucuronidase [EC:3.2.1.31] | GH2 | 5 |
| | Beta-mannosidase [EC:3.2.1.25] | GH2 | 3 |
| | Glucan 1,3-beta-glucosidase | GH16 | 1 |

| | | | |
|---|---|---|---|
| **Other Polysaccharide** | 1,3-beta-glucanase | GH16 | 1 |
| | | GH55 | 1 |
| | Alpha-L-rhamnosidase [EC:3.2.1.40] | GH78 | 1 |
| | Alpha-mannosidase [EC:3.2.1.24] (Moreira and Filho 2008) | GH38 | 4 |
| | Arabinan endo-1,5-alpha-L-arabinosidase [EC:3.2.1.99] | GH43 | 6 |
| | Endo-1,3-1,4-beta-glycanase | GH16 | 3 |
| | Endo-beta-1,3-1,4 glucanase (EC:3.2.1.73) (Kuge et al. 2015) | GH16 | 1 |
| | Mannan endo-1,4-beta-mannosidase [EC:3.2.1.78] (de Vries and Visser 2001) | GH5 | 2 |
| | Poly(beta-D-mannuronate) lyase [EC:4.2.2.3] (Zhu and Yin 2015) | PL9 | 1 |
| | Unspecified beta-glucanase [EC:3.2.1.73] | GH16 | 1 |
| | Xylanase/chitin deacetylase | CE4 | 1 |
| **Pectin** | Pectate lyase (Abbott and Boraston 2008) | PL9 | 3 |
| | Pectate lyase [EC:4.2.2.2] | PL1 | 2 |
| | Pectinesterase [EC:3.1.1.11] (Abbott and Boraston 2008) | CE8 | 1 |
| | Polygalacturonase [EC:3.2.1.15] (Abbott and Boraston 2008) | GH28 | 2 |
| | Rhamnogalacturonan endolyase [EC:4.2.2.23] | PL4 | 1 |
| | Unsaturated rhamnogalacturonyl hydrolase [EC:3.2.1.172] (Silva et al. 2016) | GH105 | 2 |
| **Pectin Accessory** | Arabinogalactan endo-1,4-beta-galactosidase [EC:3.2.1.89] (Abbott and Boraston 2008) | GH53 | 1 |
| **Sialic Acid** | Sialidase-1 [EC:3.2.1.18] (Severi et al. 2007) | GH74 | 10 |
| **Starch** | Alpha-amylase | GH13 | 20 |
| | Alpha-amylase [EC:3.2.1.1] | GH13 | 7 |
| | Alpha-amylase/alpha-mannosidase | GH57 | 1 |
| | Alpha-glucosidase | GH13 | 1 |
| | Alpha-glucosidase [EC:3.2.1.20] | GH13 | 3 |
| | | GH31 | 3 |
| | | GH97 | 5 |
| | Glucoamylase | GH15 | 2 |
| | Isoamylase | GH13 | 2 |
| | Maltodextrin glucosidase | GH13 | 1 |
| | Maltooligosyltrehalose trehalohydrolase [EC:3.2.1.141] | GH13 | 8 |
| | Oligo-1,6-glucosidase [EC:3.2.1.10] | GH13 | 3 |
| | Pullulanase [EC:3.2.1.41] | GH13 | 3 |
| **Tannan or Ferulic Acid** | Tannase and feruloyl esterase | CE1 | 1 |
| **Trehalose** | Alpha,alpha-trehalose phosphorylase [EC:2.4.1.64] | GH65 | 2 |
| | trehalose-6-phosphate hydrolase [EC:3.2.1.93] | GH13 | 1 |
| **Xylan** | Beta-1,4-xylanase [EC:3.2.1.8] (Barcelos et al. 2015) | GH5 | 1 |
| | | GH62 | 1 |

| | | | |
|---|---|---|---|
| | Cellulase/xylanase | GH10 | 2 |
| | Endo-1,4-beta-xylanase [EC:3.2.1.8] (Barcelos et al. 2015) | GH10 | 22 |
| | | GH11 | 9 |
| | | GH43 | 1 |
| | | GH62 | 1 |
| | Glucuronoarabinoxylan endo-1,4-beta-xylanase [EC:3.2.1.136] | GH30 | 4 |
| **Xylan Accessory** | Acetyl xylan esterase (Zhang et al. 2011) | CE4 | 1 |
| | Alpha-N-arabinofuranosidase [EC:3.2.1.55] (Rahman et al. 2003) | GH43 | 6 |
| | | GH51 | 17 |
| | | GH54 | 19 |
| | | GH62 | 5 |
| | Arabinoxylan arabinofuranohydrolase [EC:3.2.1.55] | GH43 | 8 |
| **Xyloglucan** | Xyloglucan-specific endo-beta-1,4-glucanase [EC:3.2.1.151] | GH12 | 2 |
| | Xyloglucan-specific exo-beta-1,4-glucanase [EC:3.2.1.155] (de Vries and Visser 2001) | GH74 | 15 |
